# Genome-wide association study of brain biochemical phenotypes reveals distinct genetic architecture of Alzheimer’s Disease related proteins

**DOI:** 10.1101/2022.05.31.493731

**Authors:** Stephanie R. Oatman, Joseph S. Reddy, Zachary Quicksall, Minerva M. Carrasquillo, Xue Wang, Chia-Chen Liu, Yu Yamazaki, Thuy T. Nguyen, Kimberly Malphrus, Michael Heckman, Kristi Biswas, Matthew Baker, Yuka A. Martens, Na Zhao, Rosa Rademakers, Michael DeTure, Melissa E. Murray, Takahisa Kanekiyo, Dennis W. Dickson, Guojun Bu, Mariet Allen, Nilüfer Ertekin-Taner

## Abstract

Alzheimer’s disease (AD) is neuropathologically characterized by amyloid-beta (Aβ) plaques and neurofibrillary tangles. Main protein components of these hallmarks include Aβ40, Aβ42, tau, phospho-tau and APOE. With the exception of the *APOE-*ε4 variant, genetic risk factors associated with brain biochemical measures of these proteins have yet to be characterized. We performed a genome-wide association study in brains of 441 AD patients for quantitative levels of these proteins collected from three distinct fractions reflecting soluble, membrane-bound and insoluble biochemical states. We identified 123 genome-wide significant associations at seven novel loci and the *APOE* locus. Genes and variants at these loci also associate with multiple AD-related measures, regulate gene expression, have cell-type specific enrichment, and roles in brain health and other neuropsychiatric diseases. Pathway analysis identified significant enrichment of shared and distinct biological pathways. Although all biochemical measures tested reflect proteins core to AD pathology, our results strongly suggest that each have unique genetic architecture and biological pathways that influence their specific biochemical states in the brain. Our novel approach of deep brain biochemical endophenotype GWAS has implications for pathophysiology of proteostasis in AD that can guide therapeutic discovery efforts focused on these proteins.

## Introduction

Alzheimer’s disease (AD) is a progressive neurodegenerative disorder, neuropathologically characterized by the accumulation of amyloid beta (Aβ) plaques and neurofibrillary tangles (NFT) in the brain^1, 2^. While AD neuropathology broadly follows characteristic patterns, heterogeneity in the composition, location and burden of the two primary lesions has been reported across post-mortem datasets^3–8^. The main component of insoluble amyloid plaques is Aβ42, while Aβ40 is often found deposited in the brain cerebrovasculature called cerebral amyloid angiopathy (CAA)^8^. Aβ is generated by the normal cleavage of the amyloid-beta precursor protein (APP), which then can oligomerize and form extracellular deposits^9, 10^. Some mutations in the *APP* gene cause a familial early-onset form of AD through modification of APP cleavage resulting in an increase in Aβ42 production^11^. Increased levels of tau are also observed in AD, along with abnormal hyperphosphorylation leading to aggregation into insoluble NFT within the cell body^2, 12^. Under normal conditions in the brain the soluble tau protein is found relatively un-phosphorylated and bound to microtubules for stabilization^13–15^. Previous genetic studies of late onset AD (LOAD) have found variants associated with the risk of developing AD; the most significant of which is the well-established *APOE-*ε4 allele^16–18^. *APOE* encodes apolipoprotein E (APOE) which functions mainly in lipid transport, but is also known to play a role in Aβ metabolism and its insoluble forms are often found co-deposited with Aβ plaques^19^. Beyond insoluble deposits of amyloid and tau species, soluble and membrane-associated biochemical states of these proteins have also been associated with AD-related phenotypes. In the temporal cortex, soluble levels of Aβ40 and Aβ42 are significantly elevated in AD compared to controls and Aβ40 levels positively correlate with disease duration^20, 21^. Membrane-associated forms of Aβ show significant positive correlation with Aβ positron emission tomography (PET) imaging in AD, while cortical Aβ42 levels have been reported to correlate with worse clinical severity and increased rate of cognitive decline^20, 21^. Moreover, it has been shown that when tau interacts with the plasma membrane, the propensity for fibrillization increases, and within the context of AD, variability in soluble tau has been shown to occur in the presence of Aβ pathology but prior to significant NFT pathology^22–31^. Apart from *APOE-*ε4, genetic risk factors associated with different brain biochemical states of distinct proteins core to the pathology of AD have yet to be identified and characterized^32^.

We hypothesize that important insights into the pathogenesis of AD may be gained by identifying genetic variants associated with variability in brain levels of AD-related protein endophenotypes including Aβ40, Aβ42, tau, phosphorylated tau (p-Tau) and APOE. Furthermore, different biochemical states (soluble, membrane, and insoluble) of AD-related proteins may have distinct genetic variants that influence their levels within the brain. Such findings may provide key insights into production or clearance pathways for these disease-associated proteins, leading to novel therapeutic targets or biomarkers. To investigate this, we utilized genetic and biochemical measures collected from the temporal cortex of 441 post-mortem AD cases. We performed a genome-wide association study (GWAS) for levels of all five proteins, collected from three biochemical states in the brain (**Figure 1**). Our findings reveal novel genetic loci and highlight the unique genetic architecture for specific biochemical states of AD-related protein endophenotypes. This study establishes deep brain biochemical endophenotype GWAS as a novel approach to dissect the biochemical heterogeneity of AD proteins which is essential to fine-tune therapeutic efforts targeting these proteins.

**Figure 1:**
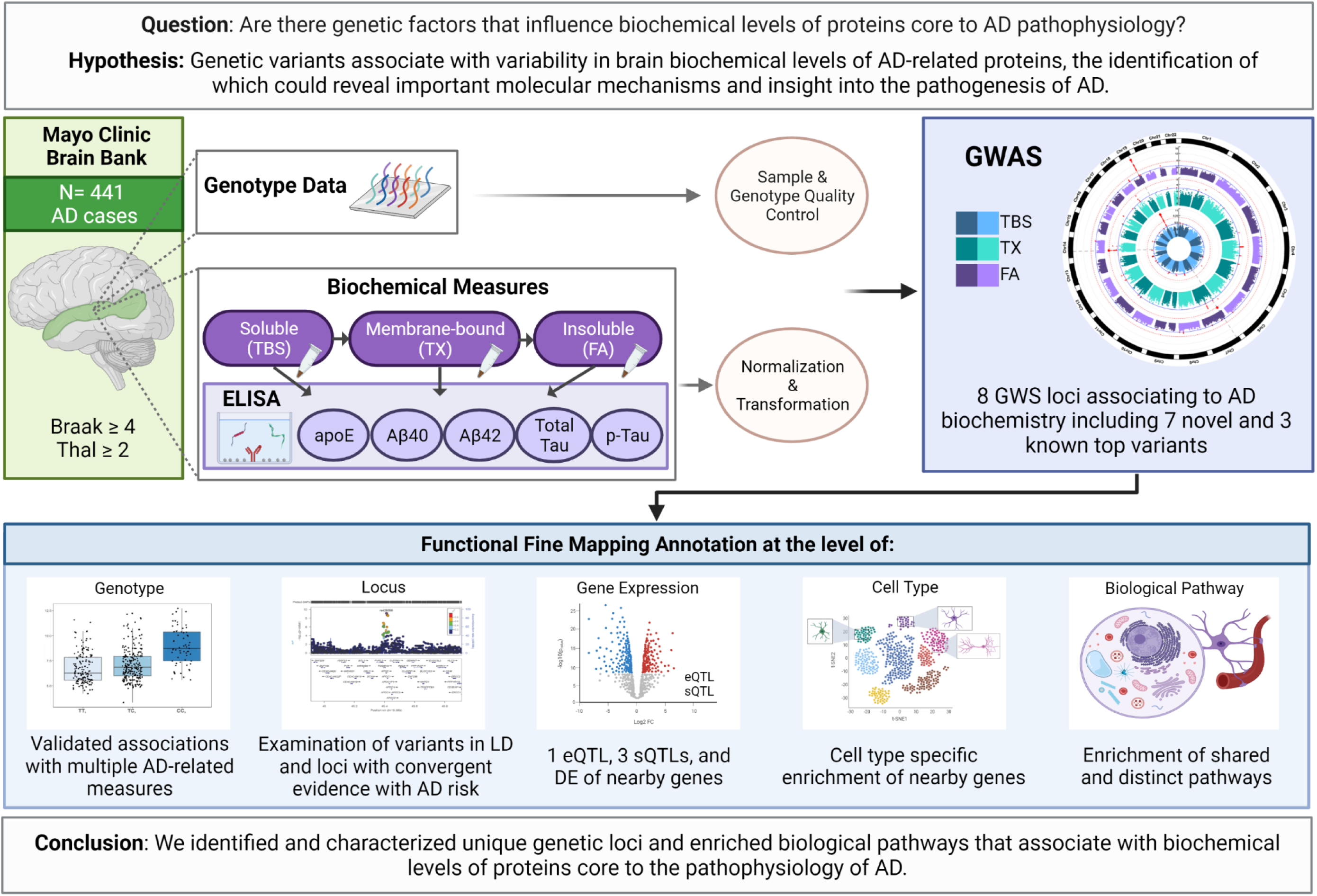
Graphical Abstract. Graphical depiction of this study. AD= Alzheimer’s Disease, N= Number, Aβ= amyloid beta, p-Tau= phosphorylated tau, TBS= Tris Buffered Saline, TX= Triton-X, FA= Formic Acid, GWS= Genome-wide significant, LD= Linkage Disequilibrium, eQTL= expression quantitative trait locus, sQTL= splicing quantitative trait locus, DE= Differential expression. Created with BioRender.com.

## Results

### Genome-wide association study identifies seven novel loci associated with AD brain biochemical endophenotypes

We utilized a cohort of 441 autopsy-confirmed AD cases from the Mayo Clinic Brain Bank with genome-wide genotypes and temporal cortex (TCX) biochemical measures of AD-related protein endophenotypes including APOE, Aβ40, Aβ42, total tau, and p-Tau from soluble (TBS), membrane (TX), and insoluble (FA) tissue fractions. Demographics including neuropathology scores are outlined in **Table S1**. Quantitative brain biochemical measures were previously collected and transformed to approximate a normal distribution (**Table S2, Figure S1**)^32^. To identify genetic associations with brain levels of AD-related proteins, genome-wide association studies were performed for each normalized biochemical fraction as well as the normalized ratio of Aβ40/42, adjusting for age, sex, the first three population principal components (PCs) in the main model, and when specified, also for *APOE* genotypes. Altogether, we identified 123 genome-wide significant (GWS, *P* < 2.89x10^-8^) SNP-endophenotype associations at 8 loci: 7 for Aβ40 (1 TBS, 5 TX, 1 FA), 4 for APOE (1 TBS, 2 TX, 1 FA), and 2 for the ratio of Aβ40/Aβ42 (1 TX, 1 FA) (**Table 1**, **Figures 2-4, Table S3**).

**Figure 2:**
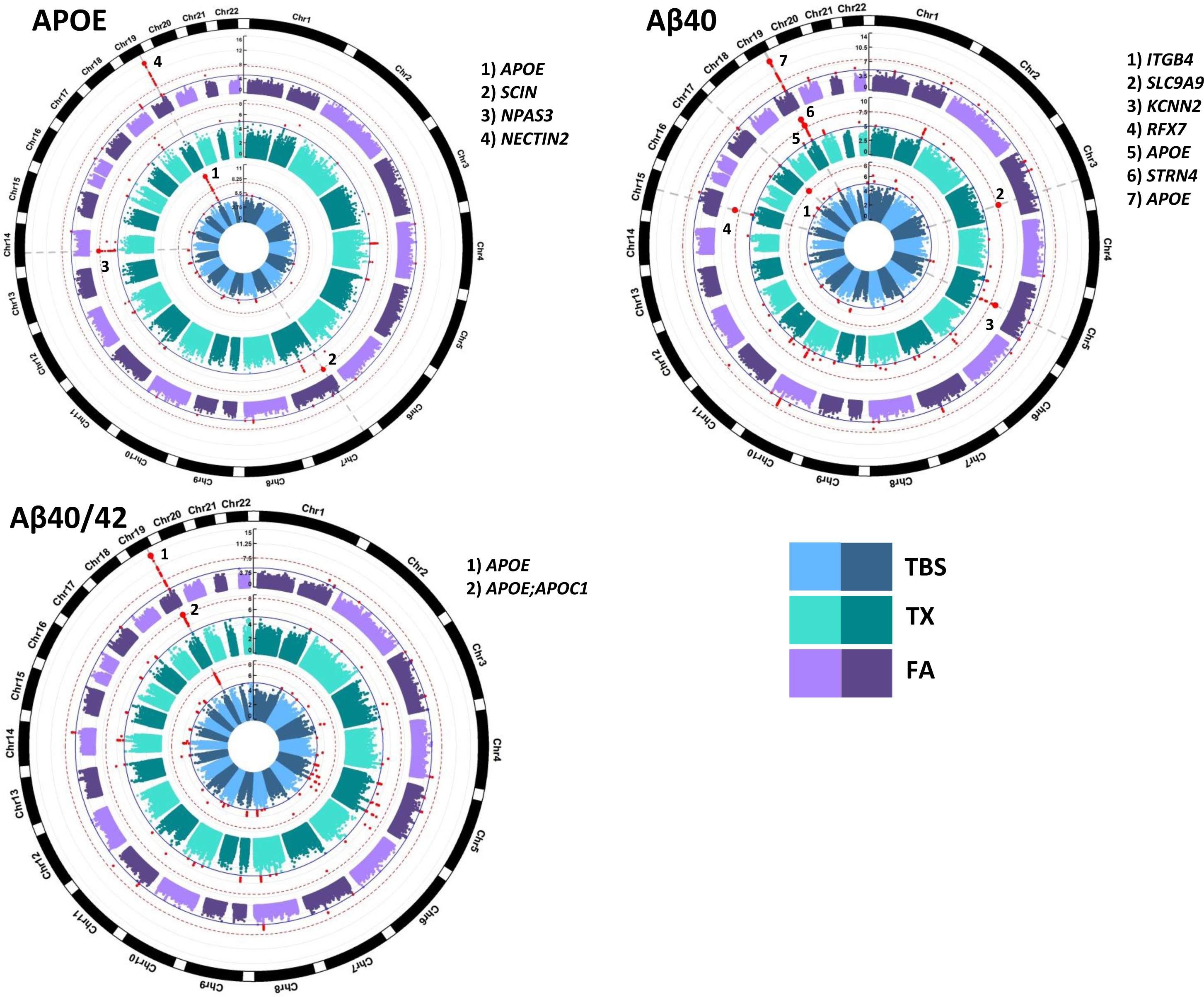
Circular Manhattan Plots of brain AD-related protein GWAS. Circular Manhattan plots for each protein measured in three biochemical fractions. Plots for proteins with SNPs that reach genome-wide significance (GWS) are shown. Red dotted line marks GWS threshold of p-value = 2.89E-08, solid blue line marks p-value = 1E-05. Top SNPs at GWS loci have dots increased in size and labeled with the closest gene name. SNPs with a p-value < 1E-05 are colored red. Radial axes measure -log10(P-value). Inner most blue circle is the soluble TBS fraction, middle green circle is the membrane TX fraction and outer most purple circle is the insoluble FA fraction.

**Figure 4:**
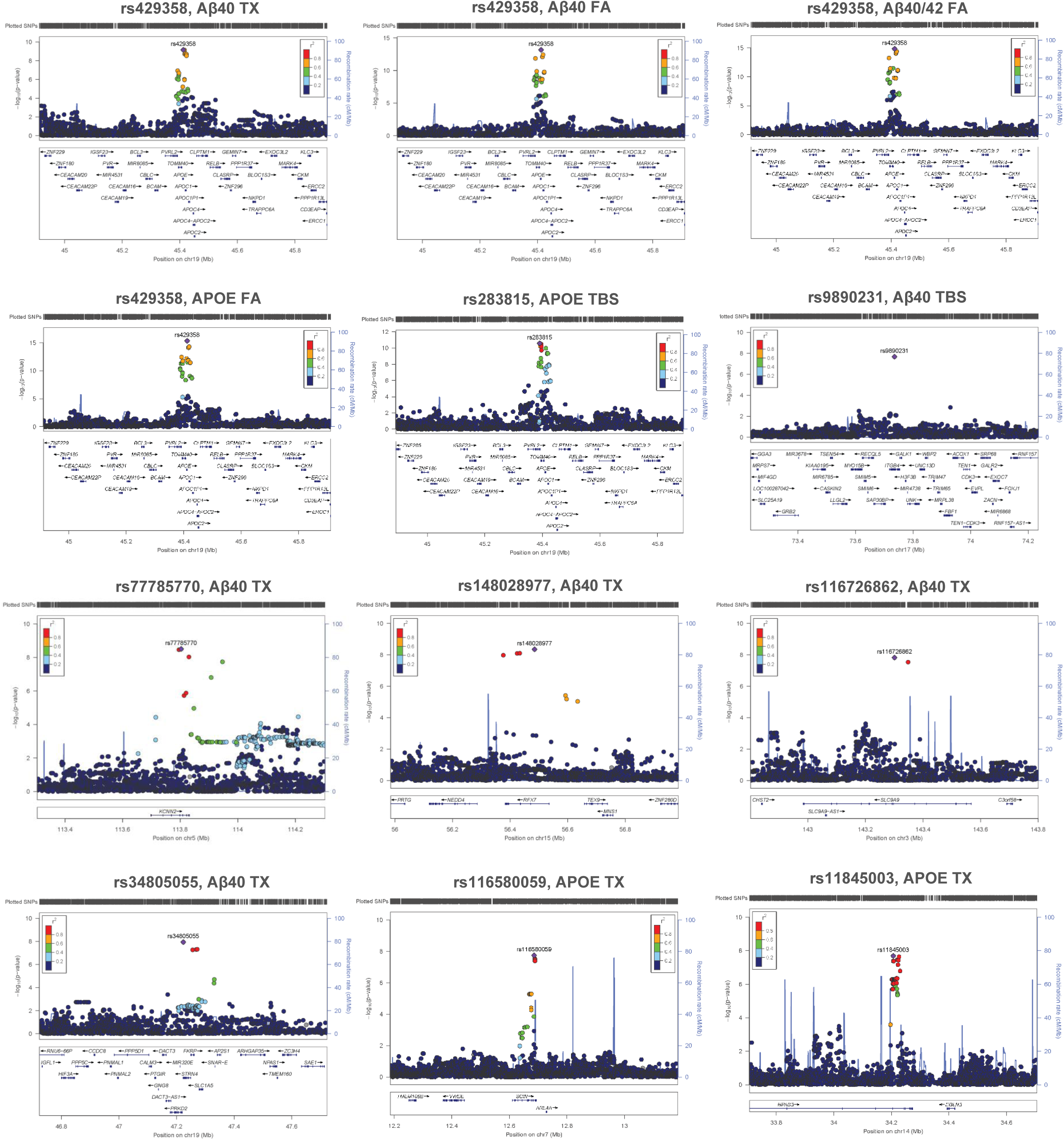
Locus Zoom Plots. Locus Zoom plots (locuszoom.org) of GWS SNPs showing associations +/- 500kb from variant of interest (labeled). Right Y-axis shows the p-value, left Y-axis shows rate of recombination, and X-axis shows position on chromosome and nearby gene positions. Each plot point represents a variant in the dataset color coded by (r^2^) value.

**Table 1:**
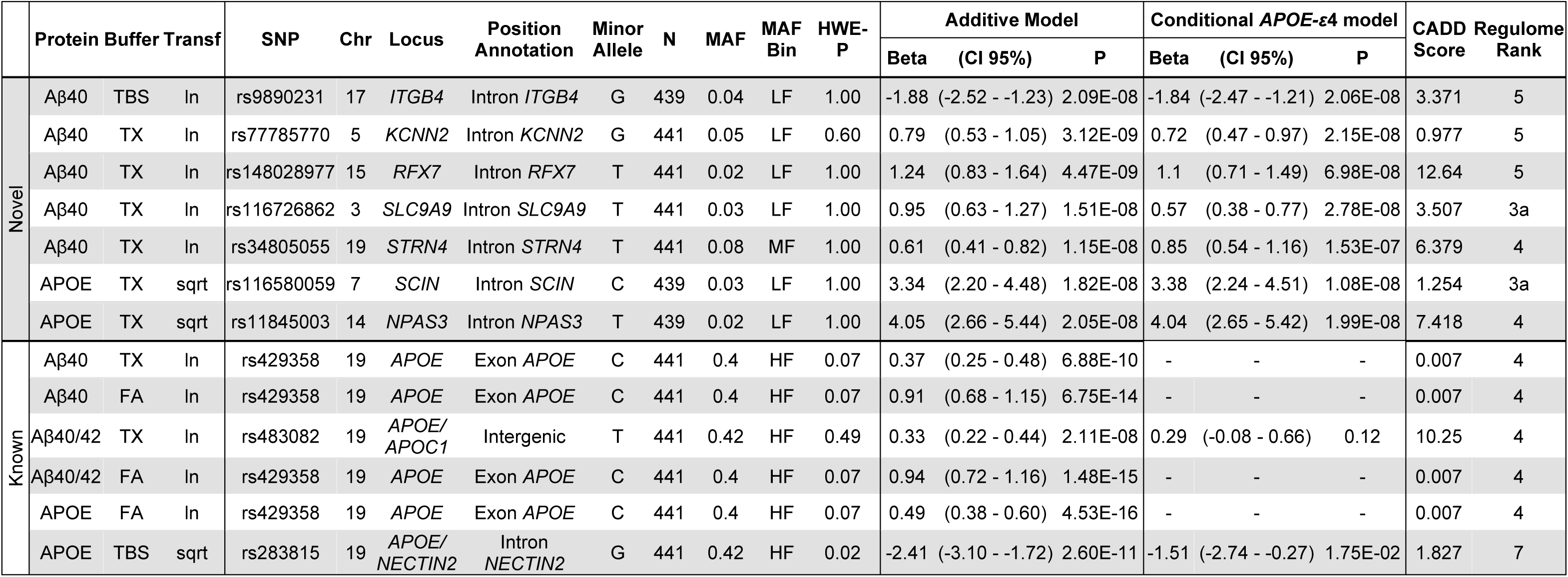
Description of genome wide significant SNPs. Descriptions of top genome-wide significant (GWS) SNPs associated with each biochemical measure in N= 441 AD cases. Seven are novel and three are known. Models include an additive model adjusted for age, sex, and PC1-3, and a conditional model conditioning on *APOE-*ε4 (rs429358) adjusted for age, sex, and PC1-3. Transf= Transformation, N= Number, MAF= Minor Allele Frequency, HWE-P= Hardy Weinberg Equilibrium P-value, CI= Confidence Interval, P= P-value.

Seven of these loci involve novel intronic variants that have not been previously implicated in genetic association studies of AD or related endophenotypes: rs116580059 near *SCIN* for APOE in TX fraction (rs116580059-APOE_TX_ (*SCIN*)), rs11845003-APOE_TX_ (*NPAS3*), rs116726862-Aβ40_TX_ (*SLC9A9*), rs148028977-Aβ40_TX_ (*RFX7*), rs34805055-Aβ40_TX_ (*STRN4*), rs77785770-Aβ40_TX_ (*KCNN2*) and rs9890231-Aβ40_TBS_ (*ITGB4*). Assessment of the index SNPs at each locus across all biochemical measures determined that each is nominally (*P* < 0.05) associated with additional biochemical measures (**Figure 3**). The estimated proportion of biochemical measure variance explained by the index SNPs based on the R^2^ of linear regression models ranged from 7% to 27%, with *APOE-ε*4 (rs429358) alone explaining between 6.6% to 14% **(Table S4)**.

**Figure 3:**
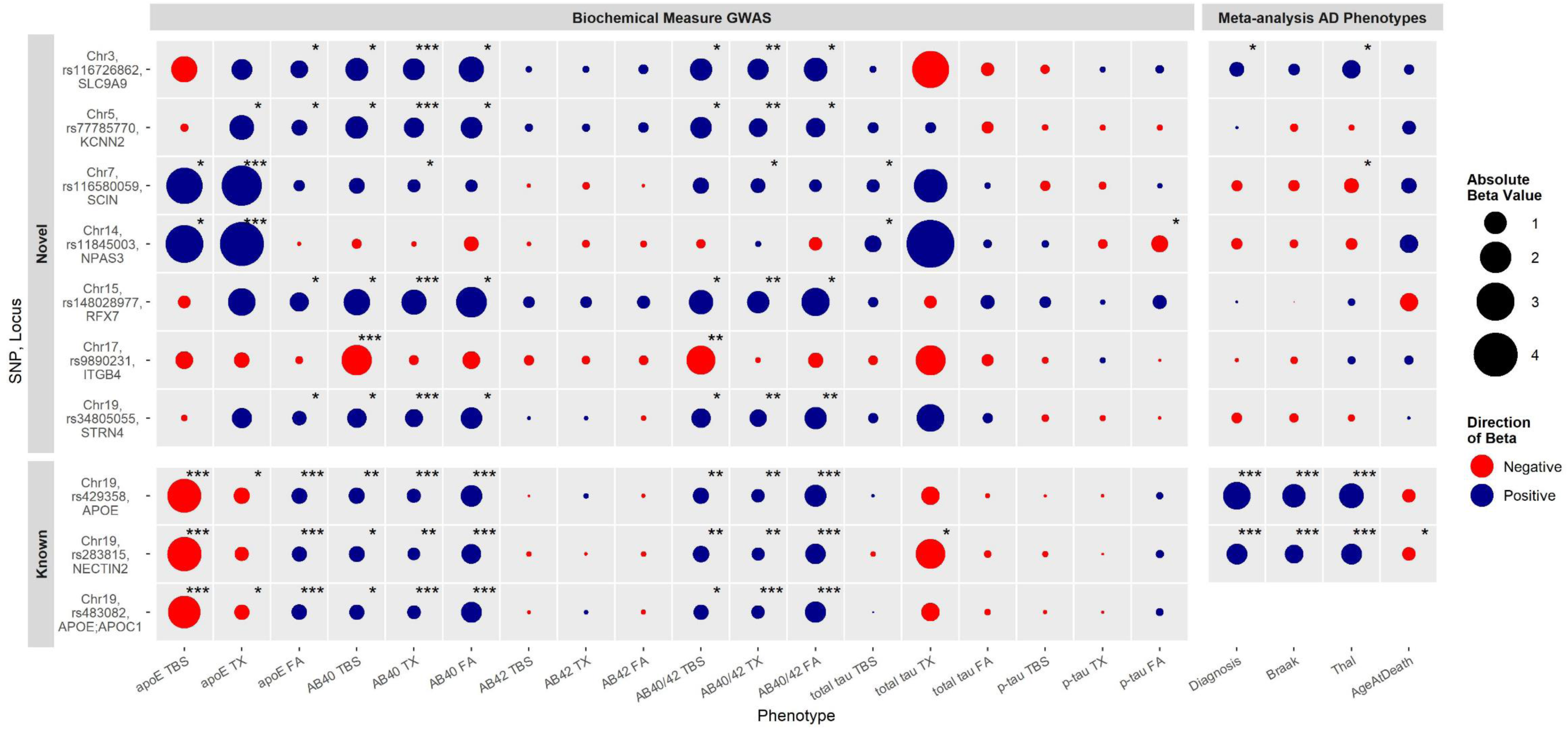
Association of Genome-wide significant SNPs across all Biochemical Measures and Meta-Analysis with AD-related Phenotypes. Associations of novel (top) and known (bottom) GWS Index SNPs across all 18 biochemical measures in the Mayo Clinic Brain Bank cohort (n=441) and results from meta-analysis with AD-related phenotypes in up to 4 cohorts (n=3,707). Meta-analysis was conducted using data from four independent autopsy datasets, namely the AMP-AD Mayo Clinic (n=344), Mount Sinai Brain Bank (MSBB, n=267), Rush (ROSMAP, n=1,091) and the Mayo expanded brain bank dataset (n=2,005). Mayo AMP-AD and expanded brain bank datasets were non-overlapping, and latter also included the 441 AD donors from the brain biochemical measures GWAS. Meta-analysis results are fixed effects models adjusted for sex and age at death when appropriate. Rs483082 was not significant after conditioning on rs429358 (*APOE-*ε4) and so was not carried forward for meta-analysis. Proxy SNPs genotypes were used for rs148028977 and rs116580059 in the Mayo expanded dataset. Note, Thal measures were only available from the expanded Mayo dataset and the AMP-AD Mayo dataset. Rs34805055 was not genotyped in the expanded Mayo dataset, therefore meta-analysis excluded this cohort for this SNP. Dot color indicates direction of beta value (blue=positive, red=negative), size of dot indicates absolute beta value. Associations with a p-value ≤ 2.89E-8 indicated by (***),1E-05 ≤ p-value < 2.89E-8 indicated by (**), and 0.05 ≤ p-value < 1E-05 indicated by (*).

More broadly, we detected 1,813 variants with a *P* < 1x10^-5^ ranging from 26 variants for total tau_TBS_ to 341 for APOE_TX_ (**Table S3**). While not reaching GWS, the most significant SNPs for the remaining traits include, Aβ42: TBS-rs147370282, TX-rs10219590, FA-rs461939; Aβ40/42: TBS-rs9890231, TX-rs483082; total tau: TBS-rs34678552, TX-rs76878089, FA-rs1634993; and p-Tau: TBS-rs2294557, TX-rs10987782, and FA-rs11651012 (**Figure S2, Table S3**). Interestingly we found associations that approach GWS for total tau_TBS_ (*P*= 8.03E-07) and total tau_TX_ (*P*= 9.48E-06) with rs117691004 which is located in the *PRKN* gene known to play a role in Parkinson’s Disease^33^ (**Table S3**).

### Multiple variants at the *APOE* locus associate with brain biochemical measures of AD-related proteins

Presence of the *APOE-*ε4 allele has previously been reported to associate with biochemical measure levels in this dataset^32^, however, using the GWAS data we can explore effects of *APOE*ε4 dose and additional genetic variation at this locus. We found a total of 30 unique GWS variants at or proximal to the *APOE* gene with at least one associated with six of the biochemical measures (**Table S3**). The known exonic AD risk *APOE-ε*4 tagging variant (rs429358) was the most significant SNP for four of the traits: Aβ40_TX_, Aβ40_FA_, Aβ40/42_FA_, APOE_FA_. The proximal *NECTIN2* intronic variant (rs283815) was the top SNP for APOE_TBS_ (**Figures 4 & 5**), and an intergenic SNP between *APOE* and *APOC1* (rs483082) was the most significant for Aβ40/42_TX_. These two SNPs are in linkage disequilibrium (LD) with *APOE-ε*4 in our dataset (rs283815: r^2^= 0.73, D’= 0.87; rs483082: r^2^= 0.92, D’= 0.99) which is likewise associated with the same biochemical traits (**Figure 3**, **Table S3**). In an *APOE-ε*4 conditional analysis, only rs283815 remains nominally significant (**Table 1**). This suggests the rs283815-APOE_TBS_ association is not fully explained by the *APOE-*ε4 allele and that multiple genetic variants at this locus may contribute to soluble APOE levels in the TCX. The rs283815 variant has previously been implicated in AD risk in males^34^ and imaging of cerebral amyloid deposition^35^, although did not survive adjustment for *APOE-ε*4 in the latter. These variants at the *APOE* region (rs429358, rs283815 and rs483082) represent the most significant associations across all fractions of Aβ40, Aβ40/42, and APOE, but no fractions of Aβ42, total tau, or p-Tau (*P* >1E-05); indicating that they likely impact disease risk through effects on APOE and Aβ40, but not tau. Furthermore, the direction of association for APOE fractions indicates a shift in biochemical state with minor allele carriers having lower soluble APOE (APOE_TBS_) and higher insoluble APOE (APOE_FA_) (**Figure 3**), suggesting a role in promoting aggregation of APOE rather than overall levels.

**Figure 5:**
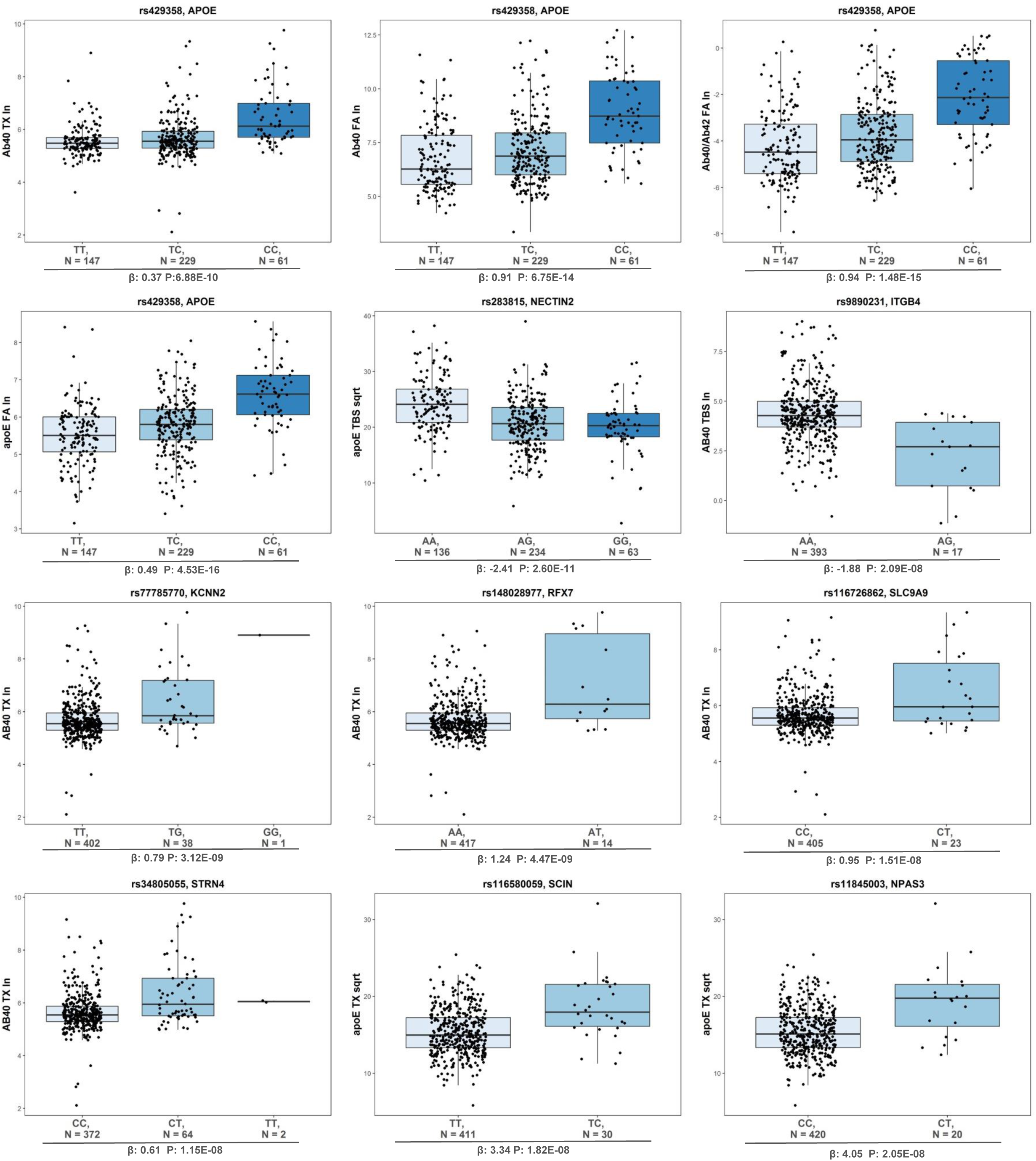
Box Plots of Genome-wide Significant SNP Genotypes. Box plots of hard-call genotypes for each genome-wide significant SNP from each biochemical measure GWAS. Each dot represents an individual sample, N = Number. Variant rs number and gene closest to GWS SNP listed at the top, beta and p-value listed at the bottom, biochemical measures on the y-axis. Genotype: Light blue-Homozygous major, Medium blue-Heterozygotes, Dark blue-Homozygous minors.

### AD brain biochemical endophenotype GWS variants also associate with disease risk, age at death, and AD-related neuropathology

To further characterize the GWS variants with respect to other AD-related phenotypes we evaluated their association with AD risk, AD-related neuropathological variables (Braak stage, Thal phase, cerebral amyloid angiopathy=CAA) and age at death in additional samples. We expanded the cohort size and validated the GWAS genotypes by genotyping these variants or their proxies in the GWAS study samples (N=441) and additional Mayo Clinic Brain Bank (MCBB) participants (N=1,564) using Taqman assays or Sanger sequencing **(Table S5)**. We refer to these 2,005 participants as the Mayo Clinic Expanded Cohort. A high level of concordance, >98%, was observed between the array-based genotyped or imputed alleles and those collected by Taqman and sequencing.

Genotypes for the index GWS variants were also extracted from three independent whole genome sequence (WGS) datasets available from the Accelerating Medicines Partnership AD (AMP-AD) study through the AD knowledge portal (www.synapse.org) which includes the Mayo Clinic RNAseq (Mayo, n=344)^36^, Mount Sinai Brain Bank (MSBB, n=267)^37^, and Rush Religious Orders Study and Memory and Aging Project (ROS-MAP, n=1,091)^38^ studies **(Table S5)**. We note that the AMP-AD Mayo and Mayo Clinic Expanded Cohort are non-overlapping. Meta-analyses for available genotypes and AD-related phenotypes were conducted across these 4 independent datasets for each SNP using fixed and random effects models **(Figure 3, Figure S3, Table S6)**. In addition, we queried a previous GWAS of CAA^39^ to determine the association of the GWS variants in the current study with this vascular AD pathology, as CAA measures were not available for the other four datasets.

As expected, the *APOE-ε*4 variant (rs429358) and the proximal *NECTIN2* variant (rs283815) significantly associated with AD risk, Thal phase, Braak stage, and as reported previously for *APOE-ε*4, also CAA^32^. Rs283815 was also associated with age at death, although the rs283815 associations were no longer significant after adjustment for *APOE-*ε2 and *APOE-*ε4 **(Figure 3, Table S6)**. We also found rs116726862 (*SLC9A9* intron, increased Aβ40_TX_) was associated with increased AD risk (*P=* 0.07), higher Thal phase (*P=* 7.50E-03), increased CAA (p=4.70E-02), and a trend for higher Braak stage (*P=* 0.07). In addition, the *SCIN* intronic SNP rs116580059 (increased APOE _TX_) was associated with lower Thal phase (p=0.015) and a trend (*P* = 0.06) for lower Braak stage **(Figure 3, Table S6)**. Although not all associations would survive Bonferroni correction, these biologically congruent associations facilitate identification of potential molecular mechanisms connecting genetic variants, biochemical measures, and other AD phenotypes. Taken together, these results validate the array-based genotype calls by an independent assay. Importantly, they implicate at least two of the novel variants near *SLC9A9* and *SCIN* more broadly in AD risk and associated neuropathology in a direction that is consistent with the brain biochemical findings.

### Brain transcriptome analyses implicate expression dysregulation at some of the novel AD brain biochemical endophenotype loci

We hypothesized that some of the variants might function though their influence on expression or splicing of nearby genes. We performed *cis*-expression quantitative trait locus (*cis*-eQTL) analysis (SNP ± 1 Mb) in three AMP-AD transcriptome datasets collected from 6 brain regions of AD cases and controls available through the AD knowledge portal and also queried independent results from the GTEx portal (www.gtexportal.org/home/, queried 08/2020)^40, 41^. We found that rs34805055-Aβ40_TX_ (intron *STRN4*) was significantly associated with downregulation of *PRKD2* gene expression in the Mayo Clinic TCX dataset (β= - 0.29, *P*= 2.6E-03) and *RN7SL364P* in the ROS-MAP dorsolateral prefrontal cortex (DLPFC) dataset (β= -0.38, *P*= 3.0E-02). The *PRKD2* gene is located approximately 3kb downstream of rs34805055 while *RN7SL364P* is a pseudogene located within an intron of *PRKD2*. In the GTEx dataset, *PRKD2* expression and splicing QTLs were found for this variant in healthy brain cortex tissue (Normalized Effect Size (NES) = -0.46, *P*= 4.1E-05), and other tissue types. The rs283815-APOE_TBS_ (*NECTIN2* intron) variant significantly associated with *TOMM40* splicing in healthy cerebellum tissue (NES= -0.94, *P*= 3.80E-16), whilst rs9890231-Aβ40_TBS_ (intron *ITGB4*) associated with *ITGB4* splicing in several tissues including healthy tibial nerve tissue (NES= -1, *P*= 6.50-09). Altogether, these results indicate that the rs34805055-Aβ40_TX_ variant may influence gene regulation and splicing of the *PRKD2* gene rather than the index gene *STRN4*, while rs9890231-Aβ40_TBS_ and rs283815-APOE_TBS_ may influence splicing of *ITGB4* and *TOMM40*, respectively. The remaining index variants do not appear to influence gene regulation in the CNS at the bulk tissue level.

We also examined each implicated locus (variant ± 1Mb, hg19) to determine if there were differentially expressed (DE) genes between AD cases and controls. We used the AMP-AD RNAseq datasets^36–38^ to assess 267 expressed genes and found that while all loci harbored DE genes in at least 1 brain region, 3 index genes and 1 gene implicated through the above QTL analysis were consistently DE in 2 or more brain regions. *KCNN2* was downregulated in AD for 3 datasets while *RFX7, SLC9A9,* and *PRKD2* were upregulated in AD for 2 (**Table S7**). Bulk tissue profiling captures changes across multiple cell types and may miss cell-specific molecular changes so we also queried results from a published single cell RNAseq (scRNA-seq) dataset^42^. We found 67 genes at the implicated loci that were DE in at least one cell type between samples with AD pathology and those with no AD pathology, 90% of which were dysregulated in neurons **(Table S8)**. Of the index genes, *ITGB4* was upregulated in astrocytes, *NECTIN2* was downregulated in neurons, while *APOE* was upregulated in neurons and microglia, and downregulated in astrocytes in AD. Interestingly, although we focus on the AD pathology vs no pathology analyses, Mathys et al.^42^ also reported DE genes between early and late AD pathology in which we see upregulation of *PRKD2* in neurons (fold change = 0.51) in late AD. Collectively these results implicate dysregulation of *PRKD2* and *ITGB4* at the novel loci identified by this study and *APOE*, *NECTIN2* and *TOMM40* at the established Chr19q13 AD locus. Larger datasets with both genetic and single cell data will be needed to further explore whether these variants influence cell-specific gene expression changes.

### Genes at the AD brain biochemical endophenotype GWAS loci are implicated in neuronal health and disease

We hypothesized that variants at the AD brain biochemical endophenotype GWAS loci may also associate with function or disease(s) of the central nervous system (CNS). We investigated all variants present in the index genes through a gene search in the GWAS catalog (https://www.ebi.ac.uk/gwas/, queried 06/07/2021)^43^. Five of the seven novel index genes have variants that associate (*P* < 1E-05) with AD-related phenotypes (**Table S6**). These include *SLC9A9* locus associated with brain Aβ40_TX_ (i.e. *SLC9A9*-Aβ40_TX_ locus) in this study and also working memory^44^, response to cholinesterase inhibitors in AD^45^ and epistatic interactions with tau measurements^46^. *SLC9A9*-Aβ40_TX_ locus was also implicated in the neuropsychiatric disorders of autism^47, 48^ and attention deficit hyperactivity disorder^49^. Further, a complete proxy of the index variant rs116726862 at *SLC9A9*-Aβ40_TX_ locus, rs115134872 (D’=1, r^2^=1), previously associated with survival in amyotrophic lateral sclerosis^50^.

Another brain biochemical endophenotype GWAS locus with AD and other neuropsychiatric disease-related associations was *NPAS3*-APOE_TX_ which also associated with neuritic and diffuse plaque measurements^51^, CSF levels of soluble TREM2^52^, epistatic interactions with tau measurements^46^, schizophrenia^53, 54^ and bipolar^53, 55, 56^ disorder. The *KCNN2*-Aβ40_TX_ locus was also associated with age of onset for AD^57^, epistatic interactions with amyloid^46^, schizophrenia^53, 54^, bipolar^53, 55, 56^ disorder and hippocampal sclerosis^58^. Additionally, *RFX7*-Aβ40_TX_ and *SCIN*-APOE_TX_ loci have associations with regional brain volume^59^ and inflammation markers^60^, respectively.

This convergent evidence supports important roles for most of the AD brain biochemical endophenotype GWAS genes and loci in neuronal health and disease.

### Gene set enrichment analysis identifies shared and distinct pathways among GWAS genes for different AD brain biochemical measures

To identify pathways that are enriched for each AD brain biochemical fraction GWAS we performed gene set enrichment analysis using GSA-SNP2^61^ with Gene Ontology (GO) terms^61^. We identified both shared and distinct significantly enriched pathway terms for each biochemical measure GWAS (**Figure 6**). Shared biological pathway terms have known roles in AD such as synapse organization, cell to cell adhesion, and immune related processes. Distinct enrichment for different biochemical fractions (TBS, TX, FA) was related to known functions for each protein, indicating that discrete molecular mechanisms likely influence specific biochemical states of each AD-related protein. For example, APOE is known to function in lipid metabolism and we found enrichment for lipoprotein clearance pathways. Notably, for soluble Aβ42_TBS_ and p-Tau_TBS_ we found enrichment in peptide cross-linking pathways indicating a genetic influence on systems that may play a role in the transition from soluble to aggregated forms of these proteins. Further, we see enrichment of synapse organization and central nervous system neuron differentiation pathways for p-Tau_FA_. This is consistent with the well-established knowledge that NFTs comprising hyper-phosphorylated tau correlate with neuronal loss and severity in AD^62^. Interestingly, we found distinct enrichment in endocytosis regulation pathways for Aβ40/42_FA_ and sensory perception of taste pathways for Aβ42_TX_ and Aβ42_FA_. These results implicate variants proximal to genes involved in both known and novel pathways that may play a role in AD pathogenesis through impacts on specific, or multiple, AD-related proteins and their distinct biochemical states in the brain.

**Figure 6:**
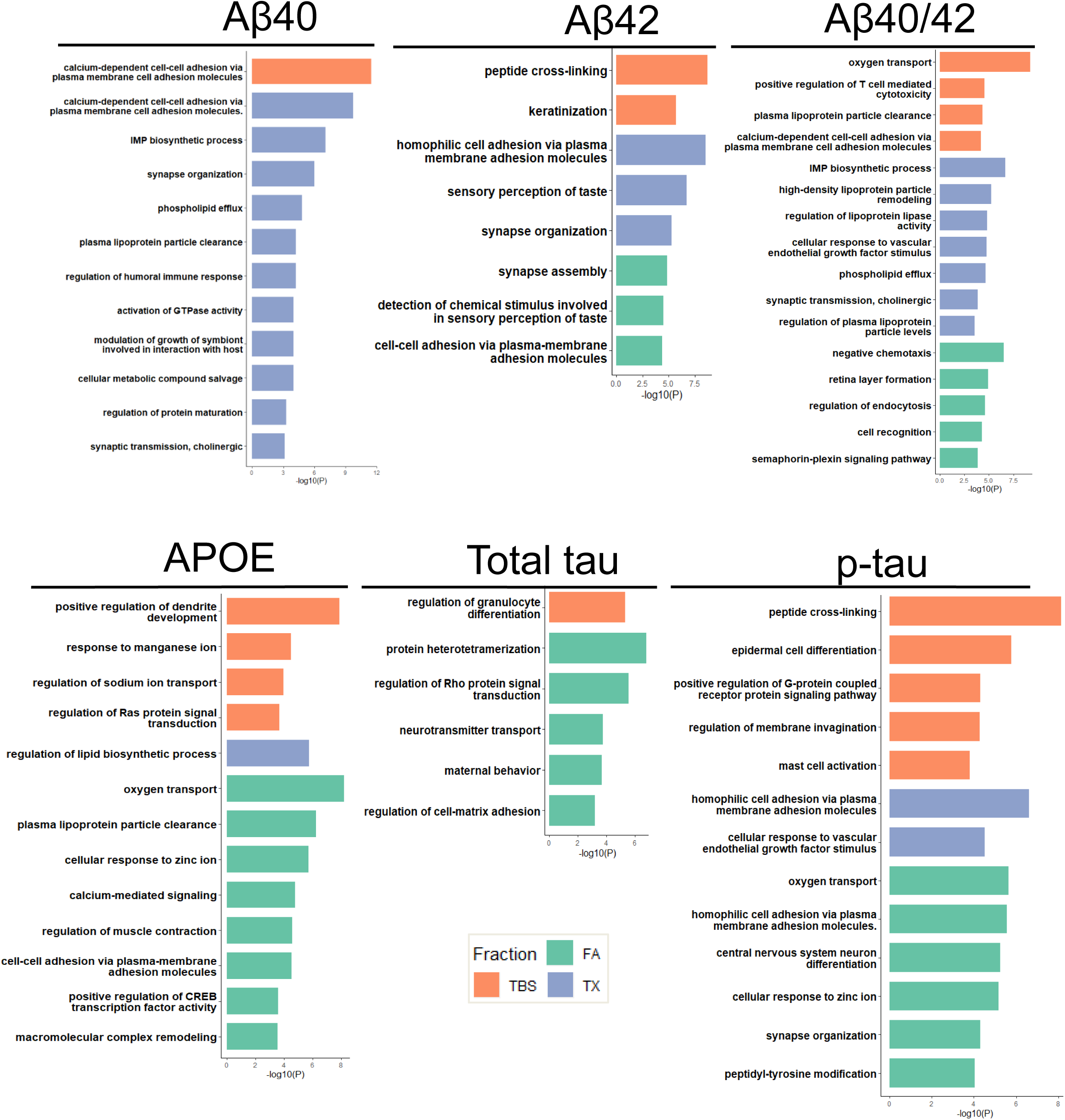
Gene set enrichment analysis in all biochemical measures. Gene set enrichment was performed via GSA-SNP2 with the MiSigDB (c5.all.v5.2). Significantly enriched pathways (q-value < 0.05) were matched with Gene Ontology (GO) IDs and input in REViGO for reduction of redundant pathways and summarization. Significant pathway groups from REViGO and the highest p-value of that group are plotted in bar plots for each protein. X-axis = -log10(P-value), Orange = soluble (TBS) fraction, Blue = membrane (TX) fraction, Green = insoluble (FA) fraction.

### Some AD risk GWAS variants also influence distinct biochemical measures

Finally, we wanted to determine if there are shared genetic risk factors for AD risk and specific brain AD-related biochemical measures. We queried large-scale AD risk GWAS^16, 17^ for the novel variants identified in this study but did not find significant association (*P* >0.05) for those outside the *APOE* locus. This would suggest that while the GWS variants may influence these brain biochemical endophenotypes, they do not have a statistically significant impact on the more heterogenous phenotype of AD risk. We also investigated 28 GWS late-onset AD (LOAD) established risk variants for association with each biochemical measure^16^. Of the 26 variants present in our dataset, 10 were nominally significant (*P* < 0.05) for at least one biochemical fraction (**Table S9**). These include *APOE-ε*4-rs429358, discussed previously, and the risk allele for rs9331896 (C) at the *CLU* locus which associates with higher brain APOE_TBS_, APOE_TX_ and total tau_TBS_, and lower p-Tau_TX_. The risk allele for rs73223431 (T) at the *PTK2B* locus associates with increased p-Tau_TX_ and p-Tau_FA_, but not APOE or Aβ measures. The protective allele of rs1080826 (A) at the *EPHA1* locus associated with lower levels of Aβ42_TX_, Aβ42_FA_ and total tau_FA_ (**Table S9**). While associations outside the *APOE* locus would not survive Bonferroni correction for 26 tests, these results using unique biochemical measures from brain tissue can still provide indications as to the underlying pathological mechanisms by which these established AD-risk variants might influence disease.

## Discussion

Genetic, model system and neuropathology studies have clearly established Aβ, tau and APOE as disease hallmarks, biomarkers and therapeutic targets in AD and other neurodegenerative diseases^63–65^. Even though there are several strategies targeting these molecules for therapeutic benefit in AD, there are critical knowledge gaps that hinder progress. All three molecules undergo complex processing and exist in heterogeneous biochemical states in the human brain. Discovering genetic and other factors that contribute to this molecular complexity and biochemical heterogeneity can yield novel therapeutic avenues. Further, the relationship of the various biochemical states of these molecules and their genetic determinants with other AD-related outcomes can help clarify the beneficial vs. detrimental mechanism of action for targeted therapies. Finally, a comprehensive genetic map of the various biochemical states of key AD proteins can help pave the way for personalized medicine targeting specific perturbed pathways in an individualized fashion.

In this study we sought to identify the genetic determinants that contribute to variability in the brain biochemical states of key AD related proteins. We report the identification of eight independent GWS loci that associate with brain levels of five hallmark AD-related proteins isolated from three tissue fractions representing soluble (TBS), membrane-bound (TX) and insoluble (FA) states of Aβ, tau and APOE. Seven of the loci are novel and associate with Aβ40, APOE, and Aβ40/Aβ42 biochemical levels. Aside from these novel loci, we also observe significant associations across the biochemical measures for the *APOE-*ε4 tagging variant (rs429358). Notably, we detected a signal within *NECTIN2* nearby the *APOE* locus that cannot be entirely explained by *APOE-ε*4. These results demonstrate the contribution of genetic factors besides *APOE-ε*4 at the *APOE* locus to the variability of brain biochemical states of AD-related proteins.

Our study also provides insights into the pathological mechanisms through which the *APOE* and novel loci may act through their influence on brain biochemical measures. The *APOE-ε*4 variant (rs429358) and the *NECTIN2* variant (rs283815) associated with measures of APOE, Aβ40, and Aβ40/42 ratio. In contrast, these variants did not show significant associations with Aβ42, total tau, and p-Tau. The results that the *APOE* locus variants were not associated with temporal cortex levels of tau proteins is consistent with previous findings in this dataset^32^. These results suggest that the *APOE* locus likely influences the risk of AD through mechanisms associated with levels of APOE and Aβ40 proteins rather than Aβ42 or tau.

Of the novel loci, *SLC9A9* which significantly associates with brain Aβ40_TX_ also has nominal associations with higher levels of many AD-related phenotypes including CAA, Braak, Thal and AD risk. In a previous study, Aβ40_TX_ levels were shown to positively correlate with CAA scores^32^. Our findings suggest that the *SLC9A9* locus may influence AD neuropathologies, including CAA, by mediating brain Aβ40_TX_ levels. *SLC9A9* encodes a sodium hydrogen exchanger with multiple functions in regulation of the endosome, an organelle that is critical for the processing of amyloid^66^. Notably, the *SLC9A9* locus has associations with other AD-related phenotypes such as working memory^44^, response to cholinesterase inhibitors in AD^45^, and with other neuropsychiatric diseases including autism, attention deficit-hyperactivity disorder, and multiple sclerosis^67–69^. Based on our findings we postulate that fundamental functions of SLC9A9 in the endosome, including amyloid processing, may underlie its influence on AD and other neuropsychiatric disease related outcomes.

Many of the other index genes discovered in our brain biochemical endophenotype GWAS have established roles relevant to AD pathology, neurological disorders, and brain function. The *KCNN2*, *RFX7*, *STRN4* and *ITGB4* loci significantly associate with brain Aβ40 levels, the first three for the membrane-bound (TX) and last for the soluble (TBS) fraction. Variants near *NPAS3* and *SCIN* associate with brain APOE_TX_ levels. *KCNN2* which encodes a calcium-activated potassium channel resides at a locus that has many other AD-related and neuropsychiatric associations (**Table S6**). In a transcriptional network analysis, *KCNN2* was found to be the top ranked network driver gene for classifying AD cases vs. controls^70^ and has been shown to have alternative splicing in AD^71^. *STRN4*, like *SLC9A9* and *KCNN2*, encodes a membrane-bound protein. STRN4 was reported to be a key binding partner and possible regulator of MAP4K^72^, the inhibition of which was shown to be neuroprotective^73, 74^. MAP4K is an upstream regulator of YAP^72^ the deficiency of which by Aβ sequestering led to neuronal necrosis in early stages of AD^75^. These findings imply that STRN4 may be a potential regulator of a molecular cascade including MAP4K and YAP involved in neuronal health and Aβ metabolism. We also found evidence through transcriptome studies that another gene at the *STRN4* locus, *PRKD2*-a protein kinase-may be the index gene. Future studies are needed to distinguish the actual functional gene at this locus.

Of the four AB40_TX_ associated loci genes, *RFX7* is the only transcription factor. Gene based rare variant analysis of *RFX7* in the Alzheimer’s Disease Neuroimaging Initiative (ADNI) cohort showed trending significance with entorhinal cortex thickness ^76^. *ITGB4*, the index gene at the Aβ40_TBS_ locus, encodes a transmembrane integrin involved in cell-to-cell adhesion, is differentially expressed in AD^77–80^ with potential roles in the blood-brain barrier^81, 82^, schizophrenia and bipolar disorder^83^.

Though most of the novel associations were with brain Aβ40 biochemical fractions, our study also discovered two significant loci for brain APOE_TX_ levels. Of these, *NPAS3*, encoding neuronal PAS domain protein, has been implicated in neurogenesis^84^, general cognitive function^85, 86^, psychiatric disorders^87^ like schizophrenia^88^, and has protein aggregation potential^89^. *NPAS3* is also known to regulate transcriptional levels of *VGF* ^90–92^ which is a key regulator in protection against AD pathogenesis in 5xFAD mice models^70^ and a top target identified by the AMP-AD consortium (agora.adknowledgeportal.org/). Finally, *SCIN* which encodes an actin-binding protein has variants that associate with inflammation markers^60^ and rate of cognitive decline in ADs^93^. The *SCIN* locus variant associated with higher APOE_TX_ is also associated with lower Thal phase in the same cohort, suggesting that higher membrane-bound levels of APOE might have a protective role in AD. Notably, upregulated expression of *SCIN* was identified as part of a pan-neurodegenerative gene signature across AD, Lewy Body disease, and ALS-FTD^94^.

In summary, investigating the functions and other genetic associations of the index genes near biochemical endophenotype GWAS loci support the conclusion that these genes and loci have functional consequences on brain health and neuropsychiatric disease. While such genetic localizations do not definitively prove the involvement of the index genes in the tested phenotypes, they nevertheless provide new testable hypotheses. Moreover, these findings underscore the potential of brain biochemical endophenotypes in the discovery of novel AD-related genes and pathways. Indeed, using the genetic association findings from our GWAS, we detected enrichment in GO pathways known to be important in AD risk as well as novel pathways. Importantly, this analysis highlighted pathways that are shared across the various biochemical endophenotypes and those that are unique to specific AD-related proteins and their unique biochemical states. In general, shared biological pathways highlight known broad biological processes in AD such as synapse organization or immune functions. In contrast, distinct pathways pinpoint processes that may relate to specific functions or biochemical states of these proteins, such as lipid metabolism for APOE and peptide cross-linking for soluble Aβ42_TBS_ and p-Tau_TBS_, respectively. This suggests that there are genetic influences which not only affect total levels but also specific biochemical states of these AD-related proteins in the brain. These findings have implications for identifying therapeutic targets that may play a role in the transition of these proteins into pathogenic biochemical states, rather than in their overall levels.

In this study, we also investigated the functional mechanisms of the GWS loci by expression and splicing QTL, as well as bulk and single cell transcriptome analyses of our and other published data. QTL analyses suggested that *STRN4*-Aβ40_TX_ rs34805055, *ITGB4*-Aβ40_TBS_ rs9890231, and *APOE/NECTIN2*-APOE_TBS_ rs283815 may modulate brain biochemical levels by impacting expression or splicing of nearby genes*, PRKD2, ITGB4* and *TOMM40*, respectively. Differential expression of genes at each locus revealed significant bulk or cell-specific transcriptional changes in AD for *PRKD2*, *ITGB4* and *APOE/NECTIN2/TOMM40*. Additional transcriptome studies, particularly cell type specific QTLs, are needed to further characterize the putative regulatory function of these variants.

In summary our study identified novel GWS loci for brain biochemical levels of AD proteins, specifically APOE and Aβ40. It is notable that these novel loci were missed in prior studies investigating CSF levels of these proteins^95^ or overall neuropathology^58^. This is likely because these studies capture more global brain changes or represent combined biochemical states of these proteins. With the exception of *APOE-ε*4, none of the GWS variants identified here associated with all three biochemical fractions of a protein, suggesting that these loci likely reflect the genetic determinants for specific biochemical states of these proteins. Our findings highlight the potential for deep biochemical phenotyping and demonstrate that this approach can dissect the genetic loci and pathways involved in the specific biochemical states of AD-related proteins, which in turn has implications for understanding disease mechanism and therapeutic development.

Our study has many strengths including extensive biochemical measures of key AD proteins Aβ, tau and APOE from three brain tissue fractions in a large sample of neuropathologically diagnosed AD patients. We provided validation and annotations for the significant GWAS loci genes and variants by leveraging additional large-scale WGS, RNA-seq, and scRNA-seq datasets with Braak, Thal, CAA, and age at death measures. We demonstrated enriched pathways that are both shared as well as those that are AD-protein and biochemical state-specific.

Nevertheless, this study has several limitations including the biochemical measures being available from a single neuropathologically-diagnosed AD cohort of 441 individuals; to our knowledge other autopsy AD cohorts with such deep brain biochemical phenotyping are lacking. Although we have leveraged other AD-related outcome associations from these and other independent samples to validate and annotate our findings, future studies in additional samples with brain biochemical measures are needed for further replications as well as increased power. Another weakness is that some of the novel GWS variants are low frequency, although this is mitigated by our validations of all index variants using a second method. We note that our AD samples have other co-pathologies, as is commonly observed in neuropathologic AD. Thus, there is a possibility that these co-pathologies may have reduced the power of this study by introducing further heterogeneity. Despite this potential confounding, we were still able to achieve GWS for *APOE* and 7 novel loci. We note that of the biochemical endophenotypes analyzed, APOE and Aβ40 had GWS associations, but not tau or Aβ42. This may be because certain proteins in specific biochemical states may be under stronger genetic control vs. may have more precise measurements reflecting their true biological variability vs. a combination of these factors. Larger scale studies with increasing measurement precision may reveal genetic factors governing the other biochemical measures. Finally, this study was conducted on non-Hispanic white individuals of Northern European descent, making it necessary to expand it to individuals of non-European ancestry.

Our results strongly suggest that, although the biochemical measures tested reflect proteins core to the pathology of AD, there are unique genetic loci associated with and enriched biological pathways for specific brain biochemical states of these proteins. These findings are expected to dissect the pathophysiology of the biochemical state of AD and finesse therapeutic target discovery efforts focused on these proteins. More broadly, this study presents a new approach that will be applicable to other neurodegenerative diseases to uncover novel mechanisms of proteostasis.

## Abbreviations

A1: Tested Allele
Aβ: Amyloid-β
AD: Alzheimer’s disease
AMP-AD-: Accelerating Medicines Partnership- Alzheimer’s Disease
APOE: Apolipoprotein E
APP: Amyloid Precursor Protein
β: Beta value
BIC: Bioinformatics core
BM: Brodmann Area
CAA: Cerebral amyloid angiopathy
CER: Cerebellum
CI: Confidence Interval
CNS: Central nervous system
DE: Differential expression
DEG: Differentially expressed genes
DLPFC: Dorsal lateral prefrontal cortex
eQTL: expression quantitative trait locus
FA: Formic acid soluble tissue fraction
FDR: False Discovery Rate
GAC: Genome analysis core
GBR: British in England and Scotland population
GO: Gene ontology
GWAS: Genome-wide association study
GWS: Genome-wide significant
HRC: Haplotype reference consortium
HWE: Hardy Weinberg Equilibrium
LD: Linkage disequilibrium
LOAD: Late-onset Alzheimer’s Disease
MAF: Minor Allele Frequency
MSBB: Mount Sinai Brain Bank
N: Number
NES: Normalized Effect Size
NFT: Neurofibrillary tangles
P: P-value
PC: Principal Component
PHF: Paired Helical Filaments
pTau-: phosphorylated tau
QQ-plot: Quantile-quantile plot
SD: Standard Deviation
SE: Standard Error
sQTL: Splicing quantitative trait locus
ROSMAP: Religious Orders Study and Rush Memory Aging Project
scRNA-seq: Single cell RNA sequencing
sqrt(CAA): Square root transformed CAA scores
TBS: Tris-buffered saline soluble tissue fraction
TCX: Temporal cortex
Transf: Transformation
TX: Detergent (1% Triton-X) soluble tissue fraction
WGS: Whole genome sequencing

## Acknowledgements

We thank the patients and families for their participation, without whom these studies would not have been possible. We thank our colleagues at the Mayo Clinic Genome Analysis Core (GAC) and Bioinformatics Core (BIC) for their collaboration.

**AD Knowledge Portal:** AMP-AD datasets: The results published here are in whole or in part based on data obtained from the AMP-AD Knowledge Portal (doi:10.7303/syn2580853 ). **Mayo Clinic:** The Mayo RNAseq study data was led by Dr. Nilüfer Ertekin-Taner, Mayo Clinic, Jacksonville, FL as part of the multi-PI U01 AG046139 (MPIs Golde, Ertekin-Taner, Younkin, Price). Samples were provided from the following sources: The Mayo Clinic Brain Bank and Banner Sun Health Research Institute. Data collection was supported through funding by NIA grants P50 AG016574, R01 AG032990, U01 AG046139, R01 AG018023, U01 AG006576, U01 AG006786, R01 AG025711, R01 AG017216, R01 AG003949, NINDS grant R01 NS080820, CurePSP Foundation, and support from Mayo Foundation. Study data includes samples collected through the Sun Health Research Institute Brain and Body Donation Program of Sun City, Arizona. The Brain and Body Donation Program is supported by the National Institute of Neurological Disorders and Stroke (U24 NS072026 National Brain and Tissue Resource for Parkinsons Disease and Related Disorders), the National Institute on Aging (P30 AG19610 Arizona Alzheimers Disease Core Center), the Arizona Department of Health Services (contract 211002, Arizona Alzheimers Research Center), the Arizona Biomedical Research Commission (contracts 4001, 0011, 05-901 and 1001 to the Arizona Parkinson’s Disease Consortium) and the Michael J. Fox Foundation for Parkinsons Research. **MSBB:** These data were generated from postmortem brain tissue collected through the Mount Sinai VA Medical Center Brain Bank and were provided by Dr. Eric Schadt from Mount Sinai School of Medicine. **ROSMAP:** Study data were provided by the Rush Alzheimer’s Disease Center, Rush University Medical Center, Chicago. Data collection was supported through funding by NIA grants P30AG10161 (ROS), R01AG15819 (ROSMAP; genomics and RNAseq), R01AG17917 (MAP), R01AG30146, R01AG36042 (5hC methylation, ATACseq), RC2AG036547 (H3K9Ac), R01AG36836 (RNAseq), R01AG48015 (monocyte RNAseq) RF1AG57473 (single nucleus RNAseq), U01AG32984 (genomic and whole exome sequencing), U01AG46152 (ROSMAP AMP-AD, targeted proteomics), U01AG46161(TMT proteomics), U01AG61356 (whole genome sequencing, targeted proteomics, ROSMAP AMP-AD), the Illinois Department of Public Health (ROSMAP), and the Translational Genomics Research Institute (genomic). Additional phenotypic data can be requested at www.radc.rush.edu. **AGORA:** The results published here are in whole or in part based on data obtained from Agora, a platform initially developed by the NIA-funded AMP-AD consortium that shares evidence in support of AD target discovery (agora.adknowledgeportal.org/). **Mayo Clinic AD-CAA:** The Mayo Clinic AD-CAA study was led by Dr. Guojun Bu and Dr. Nilufer Ertekin-Taner at Mayo Clinic, Jacksonville, FL as part of the multi-PI RF1AG051504 (MPIs Bu and Ertekin-Taner) using samples from the Mayo Clinic Brain Bank. Data collection was supported through funding by NIA grants P50AG016574, R37AG027924, Cure Alzheimer’s Fund, and support from Mayo Foundation.

**GTEx:** The Genotype-Tissue Expression (GTEx) Project was supported by the Common Fund of the Office of the Director of the National Institutes of Health, and by NCI, NHGRI, NHLBI, NIDA, NIMH, and NINDS. The data used for the analyses described in this manuscript were obtained from the GTEx Portal between 07/2020-10/2020.

**NIAGADs:** Data for this study (Kunkle et al. 2019^16^-NG00075, Lambert et al. 2013^17^-NG00036, and Cruchaga et al. 2013^95^-NG00049) were prepared, archived, and distributed by the National Institute on Aging Alzheimer’s Disease Data Storage Site (NIAGADS) at the University of Pennsylvania (U24-AG041689), funded by the National Institute on Aging.

**IGAP:** We thank the International Genomics of Alzheimer’s Project (IGAP) for providing summary results data for these analyses. The investigators within IGAP contributed to the design and implementation of IGAP and/or provided data but did not participate in analysis or writing of this report. IGAP was made possible by the generous participation of the control subjects, the patients, and their families. The i–Select chips was funded by the French National Foundation on Alzheimer’s disease and related disorders. EADI was supported by the LABEX (laboratory of excellence program investment for the future) DISTALZ grant, Inserm, Institut Pasteur de Lille, Université de Lille 2 and the Lille University Hospital. GERAD/PERADES was supported by the Medical Research Council (Grant n° 503480), Alzheimer’s Research UK (Grant n° 503176), the Wellcome Trust (Grant n° 082604/2/07/Z) and German Federal Ministry of Education and Research (BMBF): Competence Network Dementia (CND) grant n° 01GI0102, 01GI0711, 01GI0420. CHARGE was partly supported by the NIH/NIA grant R01 AG033193 and the NIA AG081220 and AGES contract N01–AG–12100, the NHLBI grant R01 HL105756, the Icelandic Heart Association, and the Erasmus Medical Center and Erasmus University. ADGC was supported by the NIH/NIA grants: U01 AG032984, U24 AG021886, U01 AG016976, and the Alzheimer’s Association grant ADGC–10–196728.

## Author Contributions

SRO, MA, JSR, and NET wrote the manuscript; NET, SRO, MA, and JSR designed the study; SRO, JSR, MA, ZSQ, MMC, and XW performed data analysis; MH consulted on statistical methods; CL, YY, YAM, NZ, TK, and GB collected biochemical measures; DWD, MD, and MEM provided neuropathological data and tissue samples; TTN and KGM isolated DNA from tissue samples; SRO, KB, MB, RR performed and consulted on targeted genotyping and sequencing. NET oversaw the study and provided direction, funding, and resources. All authors reviewed and contributed to the manuscript.

## Online Methods

### Brain samples

Post-mortem temporal cortex samples included in this study were a part of the Mayo Clinic AD-CAA (MC-CAA) study on the AD-knowledge portal (https://adknowledgeportal.synapse.org, see data sharing). All samples had a confirmed AD neuropathological diagnosis, a Braak stage ≥ four, Thal phase ≥ three, and are from non-Hispanic White decedents of Northern European descent. In total, 441 samples had both genome-wide genotyping data and biochemical measures available for analyses. This study was approved by the appropriate Mayo Clinic Institutional Review Board.

### Neuropathology

Braak stage, Thal phase, and CAA scores were measured by the Mayo Clinic Brain Bank using previously established protocols^32, 96–99^. Intermediate Braak stages were grouped with the next lowest stage as follows, stage 3.5 is 3, 4.5 is 4, and 5.5 is 5 as detailed previously^32, 39^.

### Biochemical Measures

Biochemical measures from 441 of the 469 superior temporal cortex brain samples previously described^32^ were utilized for this study based on availability of genome-wide genotypes. Biochemical measures include five AD-related proteins (APOE, Aβ40, Aβ42, tau, and phospho-tau (Thr231)) from three tissue fractions. Briefly, supernatant fractions were collected after three sequential buffer treatments of tissue homogenate and resulting pellets: first with tris-buffered saline buffer (TBS), second with detergent (TBS/1% Triton X) buffer (TX), and finally with formic acid buffer (FA), representing soluble, lipid-membrane and insoluble biochemical fractions. Quantification of AD-related biochemical measures in each fraction was performed via ELISA and normalized against total protein quantities. All biochemical measures were transformed by either the natural log or square root to achieve an approximately normal distribution including the Aβ40/42 ratio (**Figure S1**). In the subset of 441 samples analyzed, we evaluated the association of the AD-related proteins within and among these tissue fractions through pairwise correlation and found similar results as reported previously^32^ (Data not shown).

### Genotyping

DNA was isolated from brain tissue using the AutoGen245T instrument according to manufacturer’s protocols, incubated with two μl (4mg.ml) RNAseA solution (Qiagen, Germany) and stored at -80°C until use. Genome-wide genotypes from 477 samples were previously collected^39^ using the Infinium Omni2.5 Exome8 v1.3 genotyping array and results exported using the Illumina GenomeStudio software v1.9.4. Data was formatted into PLINK (v1.9) files^100, 101^ (lgen, fam, and map) and quality control of the samples and genotypes performed, as described in detail elsewhere^39^. Four hundred sixty samples passed quality control (QC), of which 441 also had biochemical measures^32^. Variants passing quality control (N= 1,383,987) were imputed to the haplotype reference consortium (HRC) panel^102, 103^ and those with an imputation quality R^2^ ≥ 0.7 and MAF ≥ 2% were kept, yielding a total of 6,726,078 variants for analysis. Genotype dosages were converted to hard calls when needed with uncertainty >0.1 set to missing. Minor allele frequencies and Hardy-Weinberg p-values were calculated for all reported variants using dosages in PLINK^100, 101^.

Genotypes for key variants, or their proxies (r^2^ = 1, D’ = 1 in 1000 Genomes EUR), were validated by Taqman genotyping or Sanger sequencing following manufacturer’s protocols. These assays were also used to collect genotypes from an additional 1,564 Mayo Clinic Brain Bank (MCBB) samples with available DNA to enable assessment with AD-related neuropathology measures of Braak stage, Thal phase, neuropathological diagnosis of AD, and age at death (combined N = 2,005, **Table S5**). These combined 2,005 AD samples are collectively referred to as the Mayo Clinic Brain Bank Expansion Cohort and are non-overlapping with the AMP-AD Mayo Clinic cohort described below. Taqman genotyping assays were performed using 10ng of dried down DNA and the QuantStudio 7 Flex system (Thermo Fisher Scientific, USA) for 7 variants (**Table S10**). Genotypes for *APOE*-rs429358 were previously collected using Taqman assays and queried from a database. One variant, *STRN4*-rs34805055, failed genotyping assay design and had no viable proxies, so Sanger sequencing was performed to validate all minor allele carrier samples. Sequencing was done on the ABI 3730 Genetic Analyzer instrument (Thermo Fisher Scientific, USA) following PCR amplification with the following primer pair: forward (5’- GGAAAGCAGCTCTGATAC) and reverse (5’- CGCATTCTGAGTCTCTG) (Integrated DNA Technologies, USA).

### AMP-AD datasets

The Mayo RNAseq study^36^, The Mount Sinai Brain Bank (MSBB) study^37^ and The Religious Orders Study and Memory and Aging Project (ROSMAP) Study^38^, were obtained from the AD-knowledge portal (https://adknowledgeportal.synapse.org). Available brain tissue RNAseq data, whole-genome genotypes, and neuropathological variables collected from these three studies were downloaded and used for fine-mapping of GWS loci and association analyses. The RNA-seq data consists of seven datasets, two from Mayo Clinic (TCX and CER), four from MSBB (BM10, BM22, BM36, and BM44), and one from ROS-MAP (DLPFC) and previously underwent consensus reprocessing (AMP-AD, RNAseq Harmonization Study)^104^. Additional QC and diagnosis harmonization of these datasets based on neuropathological measures retrieved from individual metadata files is described in detail elsewhere^39^.

In all cohorts, diagnosis was determined primarily by neuropathology made by experienced neuropathologists. The following criteria were used for diagnoses: AMP-AD Mayo dataset AD patients had a Braak stage ≥ 4 while nonADs had a Braak stage ≤ 3. AMP-AD MSBB dataset AD patients had a Braak stage ≥ 4 and CERAD score ≥ 2 while nonADs had Braak stage ≤ 3 and CERAD score ≤ 1. AMP-AD ROS-MAP dataset AD patients had a Braak stage ≥ 4 and CERAD score ≤ 2 while nonADs had Braak stage ≤ 3 and CERAD score ≥ 3. Of note, MSBB and ROS-MAP used different CERAD definitions. In ROS-MAP, CERAD score (1-4) was based on semiquantitative estimates of neuritic plaque density in one or more neocortical regions following recommendations by the Consortium to Establish a Registry for Alzheimer’s Disease (CERAD) protocol^105^. In MSBB a CERAD 1 = Normal, 2 = Definite AD, 3 = probable AD, 4 = possible AD. In ROSMAP 1= Definite AD, 2 = probable AD, 3 = Possible AD, 4 = No AD.

Whole genome sequencing (WGS) data from each of the AMP-AD cohorts was processed separately using an automated pipeline at the New York Genome Center. 150bp paired-end reads were aligned to GRCh37 human reference genome using Burrows-Wheeler Aligner^106^ (BWA-MEM v0.7.08). After marking duplicates with Picard tools^107^ (v1.83) and local read alignment around indels, base quality score recalibration (BQSR) was performed using Genome Analysis Toolkit^108^ (GATK v 3.4.0). Variant calling and joint genotyping was performed using GATK’s HaplotypeCaller (GATK v3.4.0) and GenotypeGVCFs (GATK v.3.5), respectively, to generate a multisample VCF file for each dataset. Variant quality was assessed using GATK’s variant quality score recalibration (VQSR) tool. After obtaining multi-sample VQSR-ed VCFs for each individual study from the AD knowledge portal (see data sharing), genotypes were imported into PLINK^101, 109^ (v1.9) for additional sample and variant QC using an inhouse next-generation sequencing QC pipeline. Bi-allelic autosomal variants that pass VQSR FILTER, having a genotyping rate >=98% and a minor allele frequency >= 2%, and a Bonferroni adjusted HWE p-value in controls >0.05 were retained for downstream analysis. Variants within high variability regions of the genome that can lead to spurious associations were excluded. Samples with a call rate >=98%, sex concordant with clinical information as evaluated using the inbreeding coefficient of the X-chromosome (males >=0.7, females<=0.3) and a heterozygosity estimate within 3 standard deviations (SD) of mean were retained. Relatedness among samples within each cohort was evaluated using KING^110^ robust and only one sample from each pair or family of samples related to the third degree (kinship estimate >= 0.0442) were retained. Population substructure was evaluated using Eigenstrat^111, 112^ and outliers beyond 6 SD of the top 10 principal components were removed over five iterations while refitting PCs after each iteration. After performing sample and variant QC within each cohort, data from all three datasets was merged and relatedness and population substructure were re-evaluated to exclude related samples and population outliers across all three datasets. In summary, unrelated samples of relatively homogeneous non-Hispanic White ancestry that met aforementioned sample and variant QC metrics were retained for downstream analyses.

### Statistical analysis

Power calculations were performed in R (v4.0.2) with the genpwr package. For a sample size of 441, and an alpha = 5E-08, we have 80% power to detect effect sizes of 0.42 and 0.96 when the minor allele frequency (MAF) is 0.5 and 0.05, respectively. Principal component analysis with automatic outlier exclusion was performed using Eigenstrat^111, 113^, no population outliers were identified. PLINK was utilized to perform PCA without outlier exclusion to examine samples in this cohort relative to 1000G superpopulations. PCA plots were generated using the ploty_ly() package in R (v3.6) (**Figure S4**).

PLINK (v2.00a2LM) was used to perform genome-wide association tests for variant dosage associations with each biochemical measure adjusting for age, sex and the first three population principal components (PCs) and when specified, the *APOE-*ε2 and -ε4 alleles. Genomic inflation values (λ) were calculated in R (v3.6.2) for each biochemical measure with and without adjustment of *APOE-*ε2 and -ε4 alleles (**Figure S5**); there was no evidence for genomic inflation (0.97 < λ < 1.02). QQ plots were generated in R (v3.6) with the ggplot package. (**Figure S5**). A genome-wide significance (GWS) threshold was set at *P* ≤ 2.89x10^-8^ to account for inclusion of low frequency variants^114^. To determine if GWS associations were independent from the effect of the *APOE-ε*4 allele, conditional analyses were run in PLINK 2.0 implementing the --condition command in a linear regression model conditioning on the *APOE-*ε4 tagging variant rs429358 and adjusting for age, sex, and PCs1-3. LD analysis was performed using PLINK (± 1 Mb, D’≥ 0.8 and r^2^ ≥0.2). Estimated proportion of biochemical measure variance explained by the GWS index SNPs was based on the R^2^ calculated through linear regression models regressing appropriate index SNPs on each biochemical measure.

Variants were tested for association with AD neuropathology and other related measures in the AMP-AD and expanded MCBB datasets using multi-variable regression analysis. Braak stage and Thal phase were assessed with ordinal regression in R (v4.0.2), diagnosis with logistic regression in PLINK, and age at death with linear regression in PLINK (v2.00a2LM). Samples with age at death greater than 90 years in the MCBB were redacted to 90 to parallel protocols of the AMP-AD datasets. All models included sex as a covariate, and age at death, *APOE-ε*2, and *APOE-ε*4 when appropriate or specified. Meta-analysis was performed in R (v4.0.2) with the *meta* function for both fixed and random effects models.

The AMP-AD datasets were used for assessment of each locus with brain gene expression. Differential expression analysis between diagnosis (AD case or control) and normalized gene expression levels was performed using linear regression implemented in R (v3.5.2) adjusting for age at death, sex, RNA integrity number (RIN), and sequencing batch. eQTL analysis was performed by testing variant association with CQN gene expression levels in a linear mixed model using the lme4 package in R (v3.5.2) adjusting for diagnosis, sex, age at death, RIN, tissue source, and the first three PCs, with the flow cell added as a random effects variable.

### Pathway Analysis

Gene set enrichment analysis was performed for each GWAS result with GSA-SNP2 software^61^ against the MSigDb c5.all.v5.2 database^115, 116^. Options selected include European race, GRCh37(hg19) padding build, and pathway size window of 10-200. GSA-SNP2 results were matched with Gene Ontology (GO)^117, 118^ term IDs using an in-house script. Significant GO terms and p-values were input into REViGO^119^ for summarization of significantly enriched pathways. REViGO settings were as follows: medium (0.7) allowed similarity, Homo sapiens (Gene Ontology Jan 2017) database, and SimRel semantic similarity measure. Summary bar charts were created in R (v3.6.2) with ggplot by taking reduced pathway groups from the REViGO outputs and the most significant p-value of that group for each biochemical measure.

### Variant annotations

We queried existing data and results from multiple resources to further annotate key variants and investigate the implicated loci. Associations of GWS variant dosage with sqrt(CAA) were performed previously^39^ and results queried for key variants identified in this study. Cell-specific differential gene expression analysis was queried from Mathys et al. 2019^42^ between AD pathology and no pathology samples in six cell types (excitatory neurons, inhibitory neurons, microglia, oligodendrocytes, astrocytes, and oligodendrocyte precursor cells), downloaded from the supplemental material (Table S2) on June 10, 2019 and limited to genes ± 1 Mb from the GWS variants in the Ensembl hg19 build (release 103)^120^. Only genes which passed study level significance (fdr corrected p-value ≤ 0.01 and a fold change ≥ 0.25) in at least one cell type were included in our evaluation.

Summary statistics for the LOAD GWAS (Kunkle et al. 2019^16^-NG00075, Lambert et al. 2013^17^-NG00036) and CSF GWAS (Cruchaga et al. 2013^95^-NG00049) were downloaded from NIAGADs. International Genomics of Alzheimer’s Project (IGAP) is a large three-stage study based upon genome-wide association studies (GWAS) on individuals of European ancestry. In stage 1, IGAP used genotyped and imputed data on 11,480,632 single nucleotide polymorphisms (SNPs) to meta-analyze GWAS datasets consisting of 21,982 Alzheimer’s disease cases and 41,944 cognitively normal controls from four consortia: The Alzheimer Disease Genetics Consortium (ADGC); The European Alzheimer’s disease Initiative (EADI); The Cohorts for Heart and Aging Research in Genomic Epidemiology Consortium (CHARGE); and The Genetic and Environmental Risk in AD Consortium Genetic and Environmental Risk in AD/Defining Genetic, Polygenic and Environmental Risk for Alzheimer’s Disease Consortium (GERAD/PERADES). In stage 2, 11,632 SNPs were genotyped and tested for association in an independent set of 8,362 Alzheimer’s disease cases and 10,483 controls. Meta-analysis of variants selected for analysis in stage 3A (n = 11,666) or stage 3B (n = 30,511) samples brought the final sample to 35,274 clinical and autopsy-documented Alzheimer’s disease cases and 59,163 controls.

The 1000 genomes phase_3 (GBR) dataset was queried for variants in LD (± 50kb, r^2^ ≥ 0.8, D’≥ 0.8) through Ensembl and NCBI LDlink (https://ldlink.nci.nih.gov/)^121^. The Genotype-Tissue Expression (GTEx) Project v8 (https://gtexportal.org/)^40,41^ was queried for significant eQTLs and sQTLs between August and December 2020.

### Data Sharing

The data in this manuscript will be accessed via the AD Knowledge Portal. The AD Knowledge Portal is a platform for accessing data, analyses and tools generated by the Accelerating Medicines Partnership (AMP-AD) Target Discovery Program and other National Institute on Aging (NIA)-supported programs to enable open-science practices and accelerate translational learning. The data, analyses and tools are shared early in the research cycle without a publication embargo on secondary use. Data is available for general research use according to the following requirements for data access and data attribution (https://adknowledgeportal.synapse.org/DataAccess/Instructions).

## References

1. 2017 Alzheimer’s disease facts and figures. Alzheimer’s & Dementia 13, 325–373, doi:https://doi.org/10.1016/j.jalz.2017.02.001 (2017).

2. DeTure, M. A. & Dickson, D. W. The neuropathological diagnosis of Alzheimer’s disease. Molecular Neurodegeneration 14, 32, doi:10.1186/s13024-019-0333-5 (2019).

3. Lam, B., Masellis, M., Freedman, M., Stuss, D. T. & Black, S. E. Clinical, imaging, and pathological heterogeneity of the Alzheimer’s disease syndrome. Alzheimers Res Ther 5, 1–1, doi:10.1186/alzrt155 (2013).

4. Yasuhara, O., Kawamata, T., Aimi, Y., McGeer, E. G. & McGeer, P. L. Two types of dystrophic neurites in senile plaques of Alzheimer disease and elderly non-demented cases. Neurosci Lett 171, 73–76, doi:10.1016/0304-3940(94)90608-4 (1994).

5. Janocko, N. J. et al. Neuropathologically defined subtypes of Alzheimer’s disease differ significantly from neurofibrillary tangle-predominant dementia. Acta Neuropathol 124, 681–692, doi:10.1007/s00401-012-1044-y (2012).

6. Murray, M. E. et al. Differential clinicopathologic and genetic features of late-onset amnestic dementias. Acta Neuropathol 128, 411–421, doi:10.1007/s00401-014-1302-2 (2014).

7. Mehta, R. I. & Schneider, J. A. What is ’Alzheimer’s disease’? The neuropathological heterogeneity of clinically defined Alzheimer’s dementia. Curr Opin Neurol 34, 237–245, doi:10.1097/wco.0000000000000912 (2021).

8. Lau, H. H. C., Ingelsson, M. & Watts, J. C. The existence of Abeta strains and their potential for driving phenotypic heterogeneity in Alzheimer’s disease. Acta Neuropathol, doi:10.1007/s00401-020-02201-2 (2020).

9. Golde, T. E., Eckman, C. B. & Younkin, S. G. Biochemical detection of Abeta isoforms: implications for pathogenesis, diagnosis, and treatment of Alzheimer’s disease. Biochim Biophys Acta 1502, 172–187, doi:10.1016/s0925-4439(00)00043-0 (2000).

10. Masters, C. L. et al. Alzheimer’s disease. Nature Reviews Disease Primers 1, 15056, doi:10.1038/nrdp.2015.56 (2015).

11. Bi, C., Bi, S. & Li, B. Processing of Mutant β-Amyloid Precursor Protein and the Clinicopathological Features of Familial Alzheimer’s Disease. Aging Dis 10, 383–403, doi:10.14336/AD.2018.0425 (2019).

12. Iqbal, K., Liu, F., Gong, C. X. & Grundke-Iqbal, I. Tau in Alzheimer disease and related tauopathies. Curr Alzheimer Res 7, 656–664, doi:10.2174/156720510793611592 (2010).

13. Mandelkow, E. M. et al. Tau domains, phosphorylation, and interactions with microtubules. Neurobiology of Aging 16, 355–362, doi:https://doi.org/10.1016/0197-4580(95)00025-A (1995).

14. Pooler, A. M., Noble, W. & Hanger, D. P. A role for tau at the synapse in Alzheimer’s disease pathogenesis. Neuropharmacology 76, 1–8, doi:https://doi.org/10.1016/j.neuropharm.2013.09.018 (2014).

15. Guo, T., Noble, W. & Hanger, D. P. Roles of tau protein in health and disease. Acta Neuropathol 133, 665–704, doi:10.1007/s00401-017-1707-9 (2017).

16. Kunkle, B. W. et al. Genetic meta-analysis of diagnosed Alzheimer’s disease identifies new risk loci and implicates Aβ, tau, immunity and lipid processing. Nature Genetics 51, 414–430, doi:10.1038/s41588-019-0358-2 (2019).

17. Lambert, J. C. et al. Meta-analysis of 74,046 individuals identifies 11 new susceptibility loci for Alzheimer’s disease. Nat Genet 45, 1452–1458, doi:10.1038/ng.2802 (2013).

18. Jansen, I. E. et al. Genome-wide meta-analysis identifies new loci and functional pathways influencing Alzheimer’s disease risk. Nature Genetics 51, 404–413, doi:10.1038/s41588-018-0311-9 (2019).

19. Kanekiyo, T., Xu, H. & Bu, G. ApoE and Aβ in Alzheimer’s Disease: Accidental Encounters or Partners? Neuron 81, 740–754, doi:https://doi.org/10.1016/j.neuron.2014.01.045 (2014).

20. Steinerman, J. R. et al. Distinct pools of beta-amyloid in Alzheimer disease-affected brain: a clinicopathologic study. Arch Neurol 65, 906–912, doi:10.1001/archneur.65.7.906 (2008).

21. Roberts, B. R. et al. Biochemically-defined pools of amyloid-β in sporadic Alzheimer’s disease: correlation with amyloid PET. Brain 140, 1486–1498, doi:10.1093/brain/awx057 (2017).

22. Ait-Bouziad, N. et al. Discovery and characterization of stable and toxic Tau/phospholipid oligomeric complexes. Nature Communications 8, 1678, doi:10.1038/s41467-017-01575-4 (2017).

23. Gray, E. G., PAULA-BARBOSA, M. & ROHER, A. ALZHEIMER’S DISEASE: PAIRED HELICAL FILAMENTS AND CYTOMEMBRANES. Neuropathology and Applied Neurobiology 13, 91–110, doi:https://doi.org/10.1111/j.1365-2990.1987.tb00174.x (1987).

24. Ekinci, F. J. & Shea, T. B. Phosphorylation of tau alters its association with the plasma membrane. Cell Mol Neurobiol 20, 497–508, doi:10.1023/a:1007075115574 (2000).

25. Shea, T. B. Phospholipids alter tau conformation, phosphorylation, proteolysis, and association with microtubules: implication for tau function under normal and degenerative conditions. J Neurosci Res 50, 114–122, doi:10.1002/(sici)1097-4547(19971001)50:1<114::Aid-jnr12>3.0.Co;2-b (1997).

26. Jones, E. M. et al. Interaction of tau protein with model lipid membranes induces tau structural compaction and membrane disruption. Biochemistry 51, 2539–2550, doi:10.1021/bi201857v (2012).

27. Elbaum-Garfinkle, S., Ramlall, T. & Rhoades, E. The role of the lipid bilayer in tau aggregation. Biophys J 98, 2722–2730, doi:10.1016/j.bpj.2010.03.013 (2010).

28. Brunello, C. A., Merezhko, M., Uronen, R.-L. & Huttunen, H. J. Mechanisms of secretion and spreading of pathological tau protein. Cellular and Molecular Life Sciences 77, 1721–1744, doi:10.1007/s00018-019-03349-1 (2020).

29. Alafuzoff, I. et al. Staging of neurofibrillary pathology in Alzheimer’s disease: a study of the BrainNet Europe Consortium. Brain Pathol 18, 484–496, doi:10.1111/j.1750-3639.2008.00147.x (2008).

30. Koss, D. J. et al. Soluble pre-fibrillar tau and β-amyloid species emerge in early human Alzheimer’s disease and track disease progression and cognitive decline. Acta Neuropathol 132, 875–895, doi:10.1007/s00401-016-1632-3 (2016).

31. Mattsson-Carlgren, N. et al. Aβ deposition is associated with increases in soluble and phosphorylated tau that precede a positive Tau PET in Alzheimer’s disease. Science Advances 6, eaaz2387, doi:10.1126/sciadv.aaz2387 (2020).

32. Liu, C.-C. et al. Tau and apolipoprotein E modulate cerebrovascular tight junction integrity independent of cerebral amyloid angiopathy in Alzheimer’s disease. Alzheimer’s & Dementia 16, 1372–1383, doi:https://doi.org/10.1002/alz.12104 (2020).

33. Dawson, T. M. & Dawson, V. L. The role of parkin in familial and sporadic Parkinson’s disease. Mov Disord 25 **Suppl 1**, S32–S39, doi:10.1002/mds.22798 (2010).

34. Nazarian, A., Yashin, A. I. & Kulminski, A. M. Genome-wide analysis of genetic predisposition to Alzheimer’s disease and related sex disparities. Alzheimers Res Ther 11, 5, doi:10.1186/s13195-018-0458-8 (2019).

35. Yan, Q. et al. Genome-wide association study of brain amyloid deposition as measured by Pittsburgh Compound-B (PiB)-PET imaging. Mol Psychiatry 26, 309–321, doi:10.1038/s41380-018-0246-7 (2021).

36. Allen, M. et al. Human whole genome genotype and transcriptome data for Alzheimer’s and other neurodegenerative diseases. Scientific Data 3, 160089, doi:10.1038/sdata.2016.89 (2016).

37. Wang, M. et al. The Mount Sinai cohort of large-scale genomic, transcriptomic and proteomic data in Alzheimer’s disease. Scientific Data 5, 180185, doi:10.1038/sdata.2018.185 (2018).

38. De Jager, P. L. et al. A multi-omic atlas of the human frontal cortex for aging and Alzheimer’s disease research. Scientific Data 5, 180142, doi:10.1038/sdata.2018.142 (2018).

39. Reddy, J. S. et al. Genome-wide analysis identifies a novel LINC-PINT splice variant associated with vascular amyloid pathology in Alzheimer’s disease. Acta Neuropathologica Communications 9, 93, doi:10.1186/s40478-021-01199-2 (2021).

40. Aguet, F. et al. Genetic effects on gene expression across human tissues. Nature 550, 204–213, doi:10.1038/nature24277 (2017).

41. GTEx Consortium., L. a., Aguet, F. et al. A Novel Approach to High-Quality Postmortem Tissue Procurement: The GTEx Project. Biopreservation and Biobanking 13, 311–319, doi:10.1089/bio.2015.0032 (2015).

42. Mathys, H. et al. Single-cell transcriptomic analysis of Alzheimer’s disease. Nature 570, 332–337, doi:10.1038/s41586-019-1195-2 (2019).

43. Buniello, A. et al. The NHGRI-EBI GWAS Catalog of published genome-wide association studies, targeted arrays and summary statistics 2019. Nucleic Acids Res 47, D1005–d1012, doi:10.1093/nar/gky1120 (2019).

44. Donati, G., Dumontheil, I. & Meaburn, E. L. Genome-Wide Association Study of Latent Cognitive Measures in Adolescence: Genetic Overlap With Intelligence and Education. Mind Brain Educ 13, 224–233, doi:10.1111/mbe.12198 (2019).

45. Martinelli-Boneschi, F. et al. Pharmacogenomics in Alzheimer’s disease: a genome-wide association study of response to cholinesterase inhibitors. Neurobiol Aging 34, 1711.e1717–1713, doi:10.1016/j.neurobiolaging.2012.12.008 (2013).

46. Wang, H. et al. Genome-wide interaction analysis of pathological hallmarks in Alzheimer’s disease. Neurobiol Aging 93, 61–68, doi:10.1016/j.neurobiolaging.2020.04.025 (2020).

47. Grove, J. et al. Identification of common genetic risk variants for autism spectrum disorder. Nat Genet 51, 431–444, doi:10.1038/s41588-019-0344-8 (2019).

48. Matoba, N. et al. Common genetic risk variants identified in the SPARK cohort support DDHD2 as a candidate risk gene for autism. Transl Psychiatry 10, 265, doi:10.1038/s41398-020-00953-9 (2020).

49. Mick, E. et al. Family-based genome-wide association scan of attention-deficit/hyperactivity disorder. J Am Acad Child Adolesc Psychiatry 49, 898–905.e893, doi:10.1016/j.jaac.2010.02.014 (2010).

50. Fogh, I. et al. Association of a Locus in the CAMTA1 Gene With Survival in Patients With Sporadic Amyotrophic Lateral Sclerosis. JAMA Neurol 73, 812–820, doi:10.1001/jamaneurol.2016.1114 (2016).

51. Chibnik, L. B. et al. Susceptibility to neurofibrillary tangles: role of the PTPRD locus and limited pleiotropy with other neuropathologies. Mol Psychiatry 23, 1521–1529, doi:10.1038/mp.2017.20 (2018).

52. Liu, C. & Yu, J. Genome-Wide Association Studies for Cerebrospinal Fluid Soluble TREM2 in Alzheimer’s Disease. Front Aging Neurosci 11, 297–297, doi:10.3389/fnagi.2019.00297 (2019).

53. Huang, J. et al. Cross-disorder genomewide analysis of schizophrenia, bipolar disorder, and depression. Am J Psychiatry 167, 1254–1263, doi:10.1176/appi.ajp.2010.09091335 (2010).

54. Goes, F. S. et al. Genome-wide association study of schizophrenia in Ashkenazi Jews. Am J Med Genet B Neuropsychiatr Genet 168, 649–659, doi:10.1002/ajmg.b.32349 (2015).

55. Stahl, E. A. et al. Genome-wide association study identifies 30 loci associated with bipolar disorder. Nature Genetics 51, 793–803, doi:10.1038/s41588-019-0397-8 (2019).

56. Ferreira, M. A. et al. Collaborative genome-wide association analysis supports a role for ANK3 and CACNA1C in bipolar disorder. Nat Genet 40, 1056–1058, doi:10.1038/ng.209 (2008).

57. Herold, C. et al. Family-based association analyses of imputed genotypes reveal genome-wide significant association of Alzheimer’s disease with OSBPL6, PTPRG, and PDCL3. Mol Psychiatry 21, 1608–1612, doi:10.1038/mp.2015.218 (2016).

58. Beecham, G. W. et al. Genome-wide association meta-analysis of neuropathologic features of Alzheimer’s disease and related dementias. PLoS Genet 10, e1004606, doi:10.1371/journal.pgen.1004606 (2014).

59. Alliey-Rodriguez, N. et al. NRXN1 is associated with enlargement of the temporal horns of the lateral ventricles in psychosis. Translational Psychiatry 9, 230, doi:10.1038/s41398-019-0564-9 (2019).

60. Ahola-Olli, A. V. et al. Genome-wide Association Study Identifies 27 Loci Influencing Concentrations of Circulating Cytokines and Growth Factors. Am J Hum Genet 100, 40–50, doi:10.1016/j.ajhg.2016.11.007 (2017).

61. Yoon, S. et al. Efficient pathway enrichment and network analysis of GWAS summary data using GSA-SNP2. Nucleic acids research 46, e60–e60, doi:10.1093/nar/gky175 (2018).

62. Nelson, P. T. et al. Correlation of Alzheimer disease neuropathologic changes with cognitive status: a review of the literature. J Neuropathol Exp Neurol 71, 362–381, doi:10.1097/NEN.0b013e31825018f7 (2012).

63. Martens, Y. A. et al. ApoE Cascade Hypothesis in the pathogenesis of Alzheimer’s disease and related dementias. Neuron 110, 1304–1317, doi:10.1016/j.neuron.2022.03.004 (2022).

64. Tan, C. C., Zhang, X. Y., Tan, L. & Yu, J. T. Tauopathies: Mechanisms and Therapeutic Strategies. J Alzheimers Dis 61, 487–508, doi:10.3233/JAD-170187 (2018).

65. Hampel, H. et al. The Amyloid-beta Pathway in Alzheimer’s Disease. Mol Psychiatry 26, 5481–5503, doi:10.1038/s41380-021-01249-0 (2021).

66. Nixon, R. A. Amyloid precursor protein and endosomal-lysosomal dysfunction in Alzheimer’s disease: inseparable partners in a multifactorial disease. FASEB J 31, 2729–2743, doi:10.1096/fj.201700359 (2017).

67. Patak, J., Faraone, S. V. & Zhang-James, Y. Sodium hydrogen exchanger 9 NHE9 (SLC9A9) and its emerging roles in neuropsychiatric comorbidity. American Journal of Medical Genetics Part B: Neuropsychiatric Genetics 183, 289–305, doi:https://doi.org/10.1002/ajmg.b.32787 (2020).

68. Markunas, C. A. et al. Genetic variants in SLC9A9 are associated with measures of attention-deficit/hyperactivity disorder symptoms in families. Psychiatr Genet 20, 73–81, doi:10.1097/YPG.0b013e3283351209 (2010).

69. Liu, G. et al. Genetic Variants and Multiple Sclerosis Risk Gene SLC9A9 Expression in Distinct Human Brain Regions. Molecular Neurobiology 54, 6820–6826, doi:10.1007/s12035-016-0208-5 (2017).

70. Beckmann, N. D. et al. Multiscale causal networks identify VGF as a key regulator of Alzheimer’s disease. Nature communications 11, 3942–3942, doi:10.1038/s41467-020-17405-z (2020).

71. Heinzen, E. L. et al. Alternative ion channel splicing in mesial temporal lobe epilepsy and Alzheimer’s disease. Genome Biology 8, R32, doi:10.1186/gb-2007-8-3-r32 (2007).

72. Seo, G. et al. MAP4K Interactome Reveals STRN4 as a Key STRIPAK Complex Component in Hippo Pathway Regulation. Cell Rep 32, 107860–107860, doi:10.1016/j.celrep.2020.107860 (2020).

73. Bos, P. H. et al. Development of MAP4 Kinase Inhibitors as Motor Neuron-Protecting Agents. Cell Chemical Biology 26, 1703–1715.e1737, doi:https://doi.org/10.1016/j.chembiol.2019.10.005 (2019).

74. Wu, C., Watts, M. E. & Rubin, L. L. MAP4K4 Activation Mediates Motor Neuron Degeneration in Amyotrophic Lateral Sclerosis. Cell Rep 26, 1143–1156.e1145, doi:https://doi.org/10.1016/j.celrep.2019.01.019 (2019).

75. Tanaka, H. et al. YAP-dependent necrosis occurs in early stages of Alzheimer’s disease and regulates mouse model pathology. Nature Communications 11, 507, doi:10.1038/s41467-020-14353-6 (2020).

76. Kim, D. et al. Knowledge-driven binning approach for rare variant association analysis: application to neuroimaging biomarkers in Alzheimer’s disease. BMC Med Inform Decis Mak 17, 61–61, doi:10.1186/s12911-017-0454-0 (2017).

77. Chen, W.-T. et al. Spatial Transcriptomics and In Situ Sequencing to Study Alzheimer’s Disease. Cell 182, 976–991.e919, doi:https://doi.org/10.1016/j.cell.2020.06.038 (2020).

78. Wang, X.-L. & Li, L. Cell type-specific potential pathogenic genes and functional pathways in Alzheimer’s Disease. BMC Neurology 21, 381, doi:10.1186/s12883-021-02407-1 (2021).

79. Jing, Q. et al. A Comprehensive Analysis Identified Hub Genes and Associated Drugs in Alzheimer’s Disease. Biomed Res Int 2021, 8893553–8893553, doi:10.1155/2021/8893553 (2021).

80. Chowdhury, U. N., Islam, M. B., Ahmad, S. & Moni, M. A. Systems biology and bioinformatics approach to identify gene signatures, pathways and therapeutic targets of Alzheimer’s disease. Informatics in Medicine Unlocked 21, 100439, doi:https://doi.org/10.1016/j.imu.2020.100439 (2020).

81. Milner, R. & Campbell, I. L. Increased expression of the β4 and α5 integrin subunits in cerebral blood vessels of transgenic mice chronically producing the pro-inflammatory cytokines IL-6 or IFN-α in the central nervous system. Molecular and Cellular Neuroscience 33, 429–440, doi:https://doi.org/10.1016/j.mcn.2006.09.004 (2006).

82. Verkerke, M., Hol, E. M. & Middeldorp, J. Physiological and Pathological Ageing of Astrocytes in the Human Brain. Neurochemical Research 46, 2662–2675, doi:10.1007/s11064-021-03256-7 (2021).

83. O’Brien, N. L. et al. Rare variant analysis in multiply affected families, association studies and functional analysis suggest a role for the ITGΒ4 gene in schizophrenia and bipolar disorder. Schizophrenia Research 199, 181–188, doi:https://doi.org/10.1016/j.schres.2018.03.001 (2018).

84. Kamm, G. B., Pisciottano, F., Kliger, R. & Franchini, L. F. The developmental brain gene NPAS3 contains the largest number of accelerated regulatory sequences in the human genome. Mol Biol Evol 30, 1088–1102, doi:10.1093/molbev/mst023 (2013).

85. Davies, G. et al. Genetic contributions to variation in general cognitive function: a meta-analysis of genome-wide association studies in the CHARGE consortium (N=53 949). Molecular Psychiatry 20, 183–192, doi:10.1038/mp.2014.188 (2015).

86. Trampush, J. W. et al. GWAS meta-analysis reveals novel loci and genetic correlates for general cognitive function: a report from the COGENT consortium. Molecular Psychiatry 22, 336–345, doi:10.1038/mp.2016.244 (2017).

87. Wong, J. et al. Expression of NPAS3 in the human cortex and evidence of its posttranscriptional regulation by miR-17 during development, with implications for schizophrenia. Schizophr Bull 39, 396–406, doi:10.1093/schbul/sbr177 (2013).

88. Macintyre, G. et al. Association of NPAS3 exonic variation with schizophrenia. Schizophrenia Research 120, 143–149, doi:https://doi.org/10.1016/j.schres.2010.04.002 (2010).

89. Nucifora, L. G. et al. A Mutation in NPAS3 That Segregates with Schizophrenia in a Small Family Leads to Protein Aggregation. Complex Psychiatry 2, 133–144, doi:10.1159/000447358 (2016).

90. Sha, L. et al. Transcriptional regulation of neurodevelopmental and metabolic pathways by NPAS3. Molecular Psychiatry 17, 267–279, doi:10.1038/mp.2011.73 (2012).

91. Luoma, L. M. & Berry, F. B. Molecular analysis of NPAS3 functional domains and variants. BMC Molecular Biology 19, 14, doi:10.1186/s12867-018-0117-4 (2018).

92. Yang, D. et al. NPAS3 Regulates Transcription and Expression of VGF: Implications for Neurogenesis and Psychiatric Disorders. Frontiers in Molecular Neuroscience 9, doi:10.3389/fnmol.2016.00109 (2016).

93. Sherva, R. et al. Genome-wide association study of rate of cognitive decline in Alzheimer’s disease patients identifies novel genes and pathways. Alzheimer’s & Dementia 16, 1134–1145, doi:https://doi.org/10.1002/alz.12106 (2020).

94. Noori, A., Mezlini, A. M., Hyman, B. T., Serrano-Pozo, A. & Das, S. Systematic review and meta-analysis of human transcriptomics reveals neuroinflammation, deficient energy metabolism, and proteostasis failure across neurodegeneration. Neurobiology of Disease 149, 105225, doi:https://doi.org/10.1016/j.nbd.2020.105225 (2021).

95. Cruchaga, C. et al. GWAS of Cerebrospinal Fluid Tau Levels Identifies Risk Variants for Alzheimer’s Disease. Neuron 78, 256–268, doi:https://doi.org/10.1016/j.neuron.2013.02.026 (2013).

96. Braak, H. & Braak, E. Neuropathological stageing of Alzheimer-related changes. Acta Neuropathol 82, 239–259, doi:10.1007/bf00308809 (1991).

97. Thal, D. R., Rüb, U., Orantes, M. & Braak, H. Phases of A beta-deposition in the human brain and its relevance for the development of AD. Neurology 58, 1791–1800, doi:10.1212/wnl.58.12.1791 (2002).

98. Murray, M. E. et al. Neuropathologically defined subtypes of Alzheimer’s disease with distinct clinical characteristics: a retrospective study. The Lancet Neurology 10, 785–796, doi:https://doi.org/10.1016/S1474-4422(11)70156-9 (2011).

99. Murray, M. E. et al. Clinicopathologic and 11C-Pittsburgh compound B implications of Thal amyloid phase across the Alzheimer’s disease spectrum. Brain 138, 1370–1381, doi:10.1093/brain/awv050 (2015).

100. Chang, C. C. et al. Second-generation PLINK: rising to the challenge of larger and richer datasets. GigaScience 4, doi:10.1186/s13742-015-0047-8 (2015).

101. Purcell, S. et al. PLINK: A Tool Set for Whole-Genome Association and Population-Based Linkage Analyses. The American Journal of Human Genetics 81, 559–575, doi:https://doi.org/10.1086/519795 (2007).

102. Loh, P.-R. et al. Reference-based phasing using the Haplotype Reference Consortium panel. Nature Genetics 48, 1443–1448, doi:10.1038/ng.3679 (2016).

103. McCarthy, S. et al. A reference panel of 64,976 haplotypes for genotype imputation. Nature Genetics 48, 1279–1283, doi:10.1038/ng.3643 (2016).

104. Wan, Y. W. et al. Meta-Analysis of the Alzheimer’s Disease Human Brain Transcriptome and Functional Dissection in Mouse Models. Cell Rep 32, 107908, doi:10.1016/j.celrep.2020.107908 (2020).

105. Mirra, S. S. et al. The Consortium to Establish a Registry for Alzheimer’s Disease (CERAD). Part II. Standardization of the neuropathologic assessment of Alzheimer’s disease. Neurology 41, 479–486, doi:10.1212/wnl.41.4.479 (1991).

106. Li, H. Aligning sequence reads, clone sequences and assembly contigs with BWA-MEM, <https://arxiv.org/abs/1303.3997> (2013).

107. Picard, <http://broadinstitute.github.io/picard.> (

108. Van der Auwera, G. A. et al. From FastQ data to high confidence variant calls: the Genome Analysis Toolkit best practices pipeline. Curr Protoc Bioinformatics 43, 11 10 11–33, doi:10.1002/0471250953.bi1110s43 (2013).

109. Chang, C. C. et al. Second-generation PLINK: rising to the challenge of larger and richer datasets. Gigascience 4, 7, doi:10.1186/s13742-015-0047-8 (2015).

110. Manichaikul, A. et al. Robust relationship inference in genome-wide association studies. Bioinformatics 26, 2867–2873, doi:10.1093/bioinformatics/btq559 (2010).

111. Patterson, N., Price, A. L. & Reich, D. Population structure and eigenanalysis. PLoS Genet 2, e190, doi:10.1371/journal.pgen.0020190 (2006).

112. Price, A. L. et al. Principal components analysis corrects for stratification in genome-wide association studies. Nat Genet 38, 904–909, doi:10.1038/ng1847 (2006).

113. Price, A. L. et al. Principal components analysis corrects for stratification in genome-wide association studies. Nature Genetics 38, 904–909, doi:10.1038/ng1847 (2006).

114. Fadista, J., Manning, A. K., Florez, J. C. & Groop, L. The (in)famous GWAS P-value threshold revisited and updated for low-frequency variants. European Journal of Human Genetics 24, 1202–1205, doi:10.1038/ejhg.2015.269 (2016).

115. Subramanian, A. et al. Gene set enrichment analysis: A knowledge-based approach for interpreting genome-wide expression profiles. Proceedings of the National Academy of Sciences 102, 15545–15550, doi:10.1073/pnas.0506580102 (2005).

116. Liberzon, A. et al. The Molecular Signatures Database (MSigDB) hallmark gene set collection. Cell Syst 1, 417–425, doi:10.1016/j.cels.2015.12.004 (2015).

117. Ashburner, M. et al. Gene Ontology: tool for the unification of biology. Nature Genetics 25, 25–29, doi:10.1038/75556 (2000).

118. The Gene Ontology Consortium. The Gene Ontology Resource: 20 years and still GOing strong. Nucleic Acids Research 47, D330–D338, doi:10.1093/nar/gky1055 (2018).

119. Supek, F., Bošnjak, M., Škunca, N. & Šmuc, T. REVIGO Summarizes and Visualizes Long Lists of Gene Ontology Terms. PLOS ONE 6, e21800, doi:10.1371/journal.pone.0021800 (2011).

120. Howe, K. L., et al. Ensembl 2021. Nucleic Acids Research 49, D884–D891, doi:10.1093/nar/gkaa942 (2020).

121. Machiela, M. J. & Chanock, S. J. LDlink: a web-based application for exploring population-specific haplotype structure and linking correlated alleles of possible functional variants. Bioinformatics 31, 3555–3557, doi:10.1093/bioinformatics/btv402 (2015).

