## Supplemental Files 1 for "Genome-wide association study of brain biochemical phenotypes reveals distinct genetic architecture of Alzheimer’s Disease related proteins"

**Supplemental Text Figures and Tables**

**Authors:**

Stephanie R. Oatman*[1], Joseph S. Reddy*[2], Zachary Quicksall[2], Minerva M. Carrasquillo[1], Xue Wang[2], Chia-Chen Liu[1], Yu Yamazaki[1], Thuy T. Nguyen[1], Kimberly Malphrus[1], Michael Heckman[2], Kristi Biswas[1], Matthew Baker[1], Yuka A. Martens[1], Na Zhao[1], Rosa Rademakers[1,3], Michael DeTure[1], Melissa E. Murray[1], Takahisa Kanekiyo[1], Dennis W. Dickson[1], Guojun Bu[1], Mariet Allen[1], Nilüfer Ertekin-Taner[1,4, #]

**Author Affiliations:**

1) Mayo Clinic, Department of Neuroscience, Jacksonville, FL USA

2) Mayo Clinic, Department of Quantitative Health Sciences, Jacksonville, FL USA

3) VIB-UA Center for Molecular Neurology, VIB, University of Antwerp, Antwerp, Belgium

4) Mayo Clinic, Department of Neurology, Jacksonville, FL USA

*Authors contributed equally

### Corresponding Author

**Corresponding Author Contact Information:**

Nilüfer Ertekin-Taner, MD, PhD
Mayo Clinic, Departments of Neurology and Neuroscience, 4500 San Pablo Road, Birdsall 3, Jacksonville, FL 32224., Phone: 904-953-7103, FAX: 904-953-7353.

| Subset | N | N: Sex (%) | | N: *APOE*-ε4 dose (%) | | | Mean age at death (SD) | N: Braak stage (%) | | | N: Thal (%) | | | | N: Average CAA (%) | | | |
|---|---|---|---|---|---|---|---|---|---|---|---|---|---|---|---|---|---|---|
| | | Male | Female | 0 | 1 | 2 | | 4 | 5 | 6 | 2 | 3 | 4 | 5 | 0-<1 | 1- <2 | 2-<3 | 3-4 |
| **All** | 441 | 210 (48%) | 231 (52%) | 145 (33%) | 234 (53%) | 62 (14%) | 80.0 (9.1) | 68 (15%) | 137 (31%) | 236 (54%) | 3 (1%) | 32 (7%) | 36 (8%) | 370 (84%) | 283 (64%) | 124 (28%) | 29 (7%) | 5 (1%) |
| ***APOE*-ε4 carriers (E4+)** | 296 | 139 (47%) | 157 (53%) | 0 (0%) | 234 (79%) | 62 (21%) | 80.7 (8.6) | 47 (16%) | 82 (28%) | 167 (56%) | 1 (0%) | 15 (5%) | 26 (9%) | 254 (86%) | 181 (61%) | 85 (29%) | 25 (8%) | 5 (2%) |
| ***APOE*-ε4 non carriers (E4-)** | 145 | 71 (49%) | 74 (51%) | 145 (100%) | 0 (0%) | 0 (0%) | 78.7 (9.8) | 21 (14%) | 55 (38%) | 69 (48%) | 2 (1%) | 17 (12%) | 10 (7%) | 116 (80%) | 102 (70%) | 39 (27%) | 4 (3%) | 0 (0%) |

**Table S1: Description of Dataset.** Summary demographics of the dataset analyzed in all individuals and by *APOE*-ε4 carrier status.
N= Number, SD= Standard Deviation

| Type | Extraction | Transformation | N | All | | *APOE*-ε4+ | | *APOE*-ε4- | |
|---|---|---|---|---|---|---|---|---|---|
| | | | | Mean | SD | Mean | SD | Mean | SD |
| APOE | TBS | sqrt | 441 | 21.7 | 5.0 | 20.6 | 4.5 | 24.0 | 5.3 |
| APOE | TX | sqrt | 439 | 15.6 | 3.2 | 15.3 | 3.2 | 16.2 | 3.2 |
| APOE | FA | ln | 441 | 5.8 | 0.9 | 6.0 | 0.9 | 5.5 | 0.8 |
| Aβ40 | TBS | ln | 439 | 4.3 | 1.6 | 4.4 | 1.6 | 4.0 | 1.5 |
| Aβ40 | TX | ln | 441 | 5.7 | 0.8 | 5.8 | 0.9 | 5.6 | 0.6 |
| Aβ40 | FA | ln | 441 | 7.3 | 1.7 | 7.5 | 1.8 | 6.8 | 1.6 |
| Aβ42 | TBS | ln | 441 | 6.5 | 0.6 | 6.5 | 0.6 | 6.5 | 0.6 |
| Aβ42 | TX | ln | 441 | 7.2 | 0.5 | 7.2 | 0.5 | 7.2 | 0.5 |
| Aβ42 | FA | ln | 441 | 10.9 | 0.7 | 10.9 | 0.7 | 11.0 | 0.7 |
| Aβ40/42 | TBS | ln | 439 | -2.2 | 1.5 | -2.0 | 1.5 | -2.5 | 1.4 |
| Aβ40/42 | TX | ln | 441 | -1.4 | 0.8 | -1.4 | 0.9 | -1.6 | 0.7 |
| Aβ40/42 | FA | ln | 441 | -3.7 | 1.7 | -3.4 | 1.7 | -4.2 | 1.5 |
| total tau | TBS | sqrt | 441 | 1.9 | 0.8 | 1.9 | 0.8 | 1.8 | 0.9 |
| total tau | TX | sqrt | 441 | 36.1 | 11.3 | 35.6 | 10.8 | 36.9 | 12.3 |
| total tau | FA | sqrt | 438 | 3.3 | 0.9 | 3.3 | 0.9 | 3.3 | 0.9 |
| p-tau | TBS | sqrt | 441 | 3.2 | 0.8 | 3.2 | 0.8 | 3.2 | 0.8 |
| p-tau | TX | ln | 441 | 1.7 | 0.4 | 1.7 | 0.4 | 1.8 | 0.4 |
| p-tau | FA | ln | 441 | 7.2 | 1.2 | 7.2 | 1.1 | 7.1 | 1.2 |

**Table S2: Description of Biochemical Measures.** Summary statistics of all biochemical measures analyzed in all samples and stratified by *APOE*-ε4 carrier status groups. Transformations of measures done to normalize dataset for downstream analysis. N= Number, SD= Standard Deviation, sqrt= square root, ln = natural log

| Biochemical Measure | Index SNPs at GWS loci | | Index SNPs at GWS loci excluding *APOE*-ε4 | | *APOE*-ε4 only | |
|---|---|---|---|---|---|---|
| | SNPs | $R^2$ | SNPs | $R^2$ | SNPs | $R^2$ |
| APOE TBS | rs283815, rs429358 | 0.104 | rs283815 | 0.097 | rs429358 | 0.093 |
| APOE TX | rs116580059, rs11845003 | 0.123 | - | - | - | - |
| APOE FA | - | - | - | - | rs429358 | 0.134 |
| Aβ40 TBS | rs9890231 | 0.070 | - | - | - | - |
| Aβ40 TX | rs116726862, rs34805055, rs148028977, rs77785770, rs429358 | 0.269 | rs116726862, rs34805055, rs148028977, rs77785770 | 0.222 | rs429358 | 0.086 |
| Aβ40 FA | - | - | - | - | rs429358 | 0.124 |
| Aβ40/42 TX | rs483082, rs429358 | 0.071 | rs483082 | 0.071 | rs429358 | 0.066 |
| Aβ40/42 FA | - | - | - | - | rs429358 | 0.140 |

**Table S4: Estimated proportion of biochemical variance explained by GWS index SNPs.** The $R^2$ is the estimated proportion of biochemical measure variance explained by the GWS index SNPs. Proportion was estimated through linear regression models regressing index SNPs on each biochemical measure with and without *APOE*-ε4 as appropriate, and in *APOE*-ε4 only models as appropriate. Proportion of variance was estimated only for biochemical measures with GWS SNPs. SNPs= Variants included in the linear regression model, $R^2$= Estimated proportion of variances explained by SNPs.

| Dataset | | N | AD status | | N: Sex (%) | | N: *APOE-ε4* dose (%) | | | Mean age at death (SD) | N: Braak stage (%) | | | | | | | N: Thal (%) | | | | | |
|---|---|---|---|---|---|---|---|---|---|---|---|---|---|---|---|---|---|---|---|---|---|---|---|
| | | | AD | nonAD | Male | Female | 0 | 1 | 2 | | 0 | 1 | 2 | 3 | 4 | 5 | 6 | 0 | 1 | 2 | 3 | 4 | 5 |
| Mayo Brain Bank Expansion | | 2005 | 1477 (74%) | 528 (26%) | 932 (46%) | 1073 (54%) | 957 (48%) | 826 (41%) | 218 (11%) | 79.6 (8.25) | 75 (4%) | 62 (3%) | 227 (11%) | 158 (8%) | 159 (8%) | 410 (20%) | 856 (43%) | 186 (9%) | 115 (6%) | 49 (2%) | 163 (8%) | 132 (7%) | 1021 (51%) |
| AMP-AD | Mayo | 344 | 91 (26%) | 253 (74%) | 166 (48%) | 178 (52%) | 210 (61%) | 70 (20%) | 8 (2%) | 81 (8.4) | 18 (5%) | 19 (6%) | 40 (12%) | 40 (12%) | 6 (2%) | 38 (11%) | 47 (14%) | 46 (13%) | 27 (8%) | 9 (3%) | 15 (4%) | 3 (1%) | 57 (17%) |
| AMP-AD | MSBB | 267 | 158 (59%) | 61 (23%) | 90 (34%) | 177 (66%) | 111 (42%) | 50 (19%) | 3 (1%) | 83.7 (7.47) | 7 (3%) | 23 (9%) | 35 (13%) | 42 (16%) | 28 (10%) | 33 (12%) | 99 (37%) | N/A | N/A | N/A | N/A | N/A | N/A |
| AMP-AD | ROS-MAP | 1091 | 554 (51%) | 268 (25%) | 379 (35%) | 712 (65%) | 807 (74%) | 257 (24%) | 18 (2%) | 86.9 (4.35) | 13 (1%) | 68 (6%) | 98 (9%) | 271 (25%) | 350 (32%) | 280 (26%) | 11 (1%) | N/A | N/A | N/A | N/A | N/A | N/A |

**Table S5: Description of Independent AMP-AD and Mayo Expansion Datasets.** Summary statistics of the Mayo expanded brain bank dataset and the AMP-AD WGS datasets. All cohorts had neuropathologically determined diagnoses made by experienced neuropathologists and some also had clinical data. The details of these cohorts can be found on the AD Knowledge Portal (www.synapse.org) and Methods. All datasets were used to test the association of index variants with AD diagnosis, Braak, Thal and age-at-death in a meta-analysis (Fig.3 and S3). Additionally, the Mayo expanded dataset was utilized to directly genotype the genome-wide significant index variants for confirmation of array/imputed genotypes. N= Number, SD= Standard Deviation; AMP-AD = Accelerating Partnerships in Medicine AD, MSBB= Mount Sinai Brain Bank, ROS-MAP= Religious Orders Study and Rush Memory and Aging Project.

| | | | | Novel | | | | | | | Known | |
|---|---|---|---|---|---|---|---|---|---|---|---|---|
| | | | **Index Variant** | **rs9890231** | **rs77785770** | **rs148028977** | **rs116726862** | **rs34805055** | **rs116580059** | **rs11845003** | **rs429358** | **rs283815** |
| | | | **Position Annotation** | **Intron *ITGB4*** | **Intron *KCNN2*** | **Intron *RFX7*** | **Intron *SLC9A9*** | **Intron *STRN4*** | **Intron *SCIN*** | **Intron *NPAS3*** | **Exon *APOE*** | **Intron *NECTIN2*** |
| | | | **Protein** | **Aβ40 TBS** | **Aβ40 TX** | **Aβ40 TX** | **Aβ40 TX** | **Aβ40 TX** | **APOE TX** | **APOE TX** | **Aβ40 TX Aβ40 FA Aβ40/42 FA APOE FA** | **APOE TBS** |
| Variant level annotation | **Meta-Analysis Fixed Effects-** Beta (95% CI) p-value | | **AD Diagnosis** | -0.03 (-0.27, 0.22)<br>0.84 | 0.01 (-0.27, 0.29)<br>0.93 | 0.01 (-0.39, 0.41)<br>0.96 | 0.4 (0.01, 0.79)<br>0.04 | -0.2 (-0.51, 0.12)<br>0.22 | -0.21 (-0.51, 0.09)<br>0.17 | -0.23 (-0.57, 0.1)<br>0.17 | 1.53 (1.35, 1.7)<br>1.40E-66 | 0.86 (0.72, 0.99)<br>6.50E-36 |
| | | | **Braak stage** | -0.09 (-0.27, 0.1)<br>0.37 | -0.11 (-0.32, 0.1)<br>0.3 | -1.8e-03 (-0.3, 0.3)<br>0.99 | 0.24 (-0.02, 0.5)<br>0.07 | -0.14 (-0.37, 0.09)<br>0.23 | -0.23 (-0.47, 9.3e-03)<br>0.06 | -0.12 (-0.39, 0.14)<br>0.36 | 1.07 (0.96, 1.18)<br>1.80E-77 | 0.69 (0.59, 0.79)<br>1.20E-42 |
| | | | **Thal phase** | 0.11 (-0.2, 0.42)<br>0.48 | -0.05 (-0.37, 0.27)<br>0.75 | 0.09 (-0.34, 0.52)<br>0.68 | 0.66 (0.18, 1.15)<br>7.50E-03 | -0.08 (-0.8, 0.64)<br>0.83 | -0.41 (-0.74, -0.08)<br>0.015 | -0.24 (-0.61, 0.12)<br>0.19 | 1.22 (1.05, 1.38)<br>4.50E-47 | 0.85 (0.7, 1)<br>8.40E-29 |
| | | | **Age at Death** | 0.15 (-0.47, 0.76)<br>0.64 | 0.35 (-0.36, 1.06)<br>0.33 | -0.61 (-1.61, 0.38)<br>0.23 | 0.19 (-0.65, 1.02)<br>0.66 | 0.02 (-0.6, 0.63)<br>0.96 | 0.47 (-0.35, 1.28)<br>0.26 | 0.64 (-0.24, 1.53)<br>0.15 | -0.33 (-0.69, 0.03)<br>0.07 | -0.33 (-0.64, -9.8E-03)<br>0.04 |
| | | *APOE*-ε2 and -ε4 Adjusted | **AD Diagnosis** | -5.7e-3 (-0.27, 0.26)<br>0.97 | -0.05 (-0.35, 0.26)<br>0.75 | 0.02 (-0.42, 0.45)<br>0.93 | 0.5 (0.07, 0.93)<br>0.02 | -0.15 (-0.5, 0.2)<br>0.4 | -0.17 (-0.5, 0.16)<br>0.32 | -0.24 (-0.6, 0.12)<br>0.19 | - | 0.08 (-0.12, 0.28)<br>0.43 |
| | | | **Braak stage** | -0.05 (-0.25, 0.14)<br>0.58 | -0.13 (-0.35, 0.09)<br>0.23 | -0.03 (-0.33, 0.28)<br>0.87 | 0.28 (0.01, 0.56)<br>0.04 | -0.08 (-0.32, 0.16)<br>0.51 | -0.23 (-0.48, 0.02)<br>0.07 | -0.12 (-0.39, 0.15)<br>0.38 | - | 0.12 (-0.03, 0.27)<br>0.12 |
| | | | **Thal phase** | 0.12 (-0.19, 0.44)<br>0.45 | -0.11 (-0.45, 0.22)<br>0.51 | 0.11 (-0.33, 0.56)<br>0.61 | 0.77 (0.26, 1.27)<br>0.0028 | 0.06 (-0.7, 0.83)<br>0.87 | -0.38 (-0.72, -0.04)<br>0.03 | -0.18 (-0.56, 0.19)<br>0.34 | - | 0.11 (-0.12, 0.34)<br>0.37 |
| | | | **Age at Death** | 0.07 (-0.56, 0.69)<br>0.84 | 0.37 (-0.36, 1.09)<br>0.32 | -0.57 (-1.57, 0.42)<br>0.26 | 0.11 (-0.73, 0.96)<br>0.79 | -0.09 (-0.72, 0.55)<br>0.79 | 0.56 (-0.28, 1.39)<br>0.19 | 0.63 (-0.26, 1.53)<br>0.17 | - | -0.28 (-0.76, 0.2)<br>0.26 |
| | **MC-CAA dataset associations#** (N=821) | | **CAA** (adjusted for Braak and Thal) | 0.04 (5.38E-01) | 0.10 (5.08E-02) | 0.12 (1.07E-01) | 0.13 (4.70E-02) | 0.07 (6.23E-02) | 0.02 (7.44E-01) | 0.03 (7.09E-01) | 0.18 (1.66E-16) | 0.16 (6.81E-13) |
| | **Variants in LD** | | **Mayo TCX and CER WGS datasets** (+/- 1 Mb, D'≥ 0.8 and r2 ≥0.2) | NA | NA | NA | NA | NA | rs10255616 | rs113599869, rs73258992, rs58873852 | rs769449, 19:45412778 | NA |
| | | | **1000Genomes GBR dataset** (+/- 50kb, D'≥ 0.8 and r2 ≥0.8) | NA | rs78713788, rs17361210, rs115281149, rs79124698 | rs555278768, rs146343566, rs138937239, rs139981434, rs138839116, rs190337649, rs146973898, rs144861372 | rs115134572 | rs11668878, rs62134832, rs62134819 | rs148766594, rs77048498, rs79606912 | Many (50+) | rs814573, rs157592, rs12721051, rs56131196, rs4420638, rs769449, rs10414043, rs6857, rs7256200 | rs283811, rs184017, rs157581, rs157582, rs59007384 |
| | | | **Neurological associations of LD variants from GWAS catalog search** | NA | NA | NA | Survival time of sporadic ALS | NA | NA | NA | AD, blood protein levels, CAA, lipoprotein levels, Aβ and tau CSF levels | AD, CAA, Aβ42 CSF levels |
| Gene level annotation | **GWAS catalog search for any variant within or near index gene that significantly associates with neuropsychiatric phenotypes** | | | NA | AD- Age of onset, total amyloid (E), bipolar disorder, hippocampal sclerosis of aging, schizophrenia | Subcortical grey matter volume | Response to cholinesterase inhibitors in AD, working memory, total PHF-tau (E), ADHD, autism | NA | Macrophage inflammatory protein 1b levels | Neuritic and diffuse plaques, plasma Aβ40 levels, CSF sTREM2 levels, PHF-tau levels (E), NFT (E), schizophrenia, bipolar disorder caudate nucleus volume | Many AD related GWAS | Many AD related GWAS |

**Table S6: Additional Annotations of GWS loci.** Variant level annotations include meta-analysis results using a fixed-effects model adjusted for sex and age at death when appropriate as well as *APOE*-ε2 and -ε4 when specified, variant associations with CAA from the MC-CAA GWAS dataset (Reddy et al. 2021), variants in LD from the AMP-AD Mayo CER and TCX WGS dataset as well as the 1000 Genomes GBR dataset, and nominally significant associations of variants in LD with neurological related traits reported in the NHGRI-EBI GWAS Catalog. Gene level annotation was performed for index genes with a GWAS catalog search for any variant with associations to neurological phenotypes. Significant associations (p-value < 0.05) are highlighted in red.
CI= Confidence Interval, CAA= Cerebral amyloid angiopathy, GBR= British in England and Scotland population, LD= Linkage disequilibrium, AMP-AD= Accelerating Medicines Partnership- AD, CER= Cerebellum, TCX= Temporal Cortex, PHF= Paired Helical Filaments, NFT= Neurofibrillary tangles. #From Reddy et al. 2021.

| | Protein | | Aβ40 | | | | | | Aβ42 | | | | | | Aβ40/42 | | | | | |
|---|---|---|---|---|---|---|---|---|---|---|---|---|---|---|---|---|---|---|---|---|
| | Fraction | | TBS | | TX | | FA | | TBS | | TX | | FA | | TBS | | TX | | FA | |
| Locus | SNP | A1 | β | P | β | P | β | P | β | P | β | P | β | P | β | P | β | P | β | P |
| *EPHA1* | rs10808026 | A | -0.10 | 4.61E-01 | 0.01 | 8.72E-01 | -0.06 | 6.86E-01 | -0.08 | 1.26E-01 | -0.09 | 3.43E-02 | -0.14 | 2.98E-02 | -0.02 | 8.72E-01 | 0.11 | 1.61E-01 | 0.08 | 6.08E-01 |
| *INPP5D* | rs10933431 | G | 0.08 | 5.59E-01 | -0.02 | 7.38E-01 | 0.00 | 9.86E-01 | 0.01 | 8.09E-01 | -0.04 | 3.57E-01 | -0.04 | 4.80E-01 | 0.06 | 6.19E-01 | 0.02 | 8.32E-01 | 0.04 | 7.76E-01 |
| *SORL1* | rs11218343 | C | 0.20 | 5.35E-01 | 0.00 | 9.85E-01 | -0.18 | 6.17E-01 | 0.00 | 9.78E-01 | -0.04 | 6.99E-01 | -0.17 | 2.51E-01 | 0.21 | 5.06E-01 | 0.04 | 8.06E-01 | -0.01 | 9.81E-01 |
| *OARD1* | rs114812713 | C | -0.41 | 1.37E-01 | 0.03 | 8.62E-01 | -0.16 | 6.02E-01 | -0.18 | 9.34E-02 | -0.06 | 4.73E-01 | -0.13 | 3.03E-01 | -0.23 | 3.78E-01 | 0.09 | 5.53E-01 | -0.03 | 9.23E-01 |
| *NYAP1* | rs12539172 | T | 0.11 | 3.66E-01 | 0.06 | 3.35E-01 | 0.35 | 7.02E-03 | -0.01 | 8.64E-01 | 0.00 | 9.81E-01 | 0.01 | 8.10E-01 | 0.12 | 3.02E-01 | 0.06 | 3.49E-01 | 0.34 | 7.42E-03 |
| *SLC24A4* | rs12881735 | C | 0.16 | 2.04E-01 | 0.10 | 1.34E-01 | 0.09 | 5.26E-01 | 0.01 | 9.07E-01 | 0.01 | 7.73E-01 | -0.04 | 4.77E-01 | 0.16 | 1.93E-01 | 0.09 | 1.91E-01 | 0.13 | 3.38E-01 |
| *CYB561; ACE* | rs138190086 | A | -0.02 | 9.54E-01 | -0.06 | 7.75E-01 | 0.24 | 5.45E-01 | 0.03 | 8.48E-01 | 0.03 | 7.68E-01 | -0.05 | 7.58E-01 | -0.05 | 8.81E-01 | -0.09 | 6.49E-01 | 0.29 | 4.50E-01 |
| *FERMT2* | rs17125924 | G | -0.13 | 4.89E-01 | 0.14 | 1.85E-01 | 0.22 | 3.03E-01 | 0.05 | 4.93E-01 | 0.04 | 5.04E-01 | 0.07 | 4.33E-01 | -0.18 | 3.02E-01 | 0.09 | 3.60E-01 | 0.15 | 4.68E-01 |
| *MEF2C-AS1* | rs190982 | G | -0.01 | 9.47E-01 | 0.03 | 6.36E-01 | -0.07 | 5.26E-01 | 0.03 | 5.45E-01 | 0.01 | 7.93E-01 | -0.03 | 5.67E-01 | -0.03 | 7.49E-01 | 0.02 | 7.55E-01 | -0.05 | 6.83E-01 |
| *CYYR1; ADAMTS1* | rs2830500 | A | 0.07 | 5.30E-01 | 0.03 | 6.18E-01 | 0.03 | 8.27E-01 | 0.03 | 4.95E-01 | 0.02 | 6.01E-01 | -0.01 | 8.97E-01 | 0.04 | 6.90E-01 | 0.01 | 8.54E-01 | 0.03 | 7.79E-01 |
| *SPI1* | rs3740688 | G | -0.01 | 9.45E-01 | 0.02 | 7.32E-01 | -0.02 | 8.35E-01 | -0.04 | 3.97E-01 | -0.04 | 2.70E-01 | -0.05 | 2.84E-01 | 0.03 | 7.86E-01 | 0.06 | 3.23E-01 | 0.03 | 8.07E-01 |
| *ABCA7* | rs3752246 | G | -0.06 | 6.66E-01 | 0.01 | 9.04E-01 | -0.01 | 9.63E-01 | 0.00 | 9.45E-01 | -0.03 | 4.92E-01 | 0.02 | 7.48E-01 | -0.07 | 6.33E-01 | 0.04 | 6.00E-01 | -0.03 | 8.52E-01 |
| *PICALM; EED* | rs3851179 | T | 0.03 | 7.78E-01 | -0.05 | 3.84E-01 | 0.05 | 6.83E-01 | 0.02 | 6.70E-01 | 0.03 | 4.03E-01 | -0.02 | 7.56E-01 | 0.01 | 9.28E-01 | -0.08 | 1.76E-01 | 0.07 | 5.79E-01 |
| *APOE* | rs429358 | C | 0.50 | 9.02E-06 | 0.37 | 6.88E-10 | 0.91 | 6.75E-14 | -0.01 | 8.73E-01 | 0.04 | 2.67E-01 | -0.03 | 6.27E-01 | 0.51 | 1.79E-06 | 0.33 | 7.08E-08 | 0.94 | 1.48E-15 |
| *GPR141; NME8* | rs4723711 | T | -0.04 | 7.41E-01 | -0.01 | 8.89E-01 | 0.05 | 6.91E-01 | -0.02 | 7.38E-01 | -0.02 | 5.88E-01 | -0.02 | 6.96E-01 | -0.02 | 8.26E-01 | 0.01 | 8.57E-01 | 0.07 | 5.63E-01 |
| *CR1* | rs4844610 | A | 0.10 | 4.60E-01 | 0.02 | 7.77E-01 | 0.14 | 3.70E-01 | 0.02 | 7.72E-01 | -0.01 | 8.25E-01 | 0.02 | 7.50E-01 | 0.09 | 5.00E-01 | 0.03 | 6.82E-01 | 0.12 | 4.31E-01 |
| *ADAM10;LOC101928725* | rs593742 | G | -0.09 | 4.39E-01 | -0.03 | 6.07E-01 | -0.14 | 2.78E-01 | -0.08 | 9.50E-02 | -0.06 | 1.43E-01 | -0.01 | 7.97E-01 | -0.01 | 9.03E-01 | 0.02 | 7.23E-01 | -0.13 | 3.14E-01 |
| *CASS4* | rs6024870 | A | 0.06 | 7.70E-01 | -0.06 | 5.33E-01 | 0.05 | 7.99E-01 | 0.03 | 7.05E-01 | 0.01 | 8.86E-01 | 0.06 | 4.75E-01 | 0.03 | 8.61E-01 | -0.07 | 4.84E-01 | -0.01 | 9.65E-01 |
| *WWOX; MAF* | rs62039712 | A | -0.03 | 8.75E-01 | 0.09 | 4.18E-01 | 0.14 | 5.33E-01 | -0.04 | 6.07E-01 | -0.10 | 1.20E-01 | -0.16 | 8.79E-02 | 0.01 | 9.72E-01 | 0.19 | 8.64E-02 | 0.30 | 1.69E-01 |
| *BIN1; CYP27C1* | rs6733839 | T | 0.05 | 6.37E-01 | 0.03 | 6.04E-01 | 0.07 | 5.19E-01 | 0.02 | 6.41E-01 | 0.04 | 2.00E-01 | 0.02 | 6.20E-01 | 0.03 | 7.70E-01 | -0.01 | 8.08E-01 | 0.05 | 6.51E-01 |
| *IQCK* | rs7185636 | C | 0.30 | 2.75E-02 | 0.07 | 3.46E-01 | 0.09 | 5.59E-01 | 0.01 | 8.25E-01 | 0.02 | 5.80E-01 | 0.01 | 9.21E-01 | 0.29 | 2.44E-02 | 0.04 | 5.46E-01 | 0.08 | 5.75E-01 |
| *PTK2B* | rs73223431 | T | 0.10 | 3.23E-01 | 0.00 | 9.86E-01 | 0.05 | 6.79E-01 | 0.02 | 5.66E-01 | 0.05 | 1.25E-01 | 0.08 | 8.13E-02 | 0.08 | 4.04E-01 | -0.05 | 3.58E-01 | -0.04 | 7.50E-01 |
| *USP6NL; ECHDC3* | rs7920721 | G | -0.15 | 1.67E-01 | -0.05 | 3.79E-01 | -0.08 | 5.08E-01 | -0.01 | 7.51E-01 | -0.01 | 7.01E-01 | 0.06 | 1.91E-01 | -0.13 | 1.86E-01 | -0.04 | 5.21E-01 | -0.14 | 2.13E-01 |
| *MS4A2;MS4A6A* | rs7933202 | C | 0.11 | 3.67E-01 | 0.07 | 2.37E-01 | 0.12 | 3.56E-01 | -0.02 | 6.70E-01 | -0.01 | 8.08E-01 | 0.00 | 9.70E-01 | 0.13 | 2.43E-01 | 0.08 | 1.90E-01 | 0.12 | 3.50E-01 |
| *CLU* | rs9331896 | C | 0.15 | 1.74E-01 | 0.06 | 3.18E-01 | 0.01 | 9.43E-01 | -0.02 | 6.49E-01 | 0.01 | 7.53E-01 | -0.07 | 1.46E-01 | 0.17 | 1.09E-01 | 0.05 | 4.23E-01 | 0.08 | 4.86E-01 |
| *TNFRSF21; CD2AP* | rs9473117 | C | 0.07 | 5.72E-01 | 0.00 | 9.98E-01 | 0.12 | 3.71E-01 | 0.02 | 6.85E-01 | -0.04 | 2.85E-01 | -0.05 | 3.67E-01 | 0.05 | 6.39E-01 | 0.04 | 5.28E-01 | 0.17 | 1.90E-01 |

| | Protein | | APOE | | | | | | Total tau | | | | | | p-Tau | | | | | |
|---|---|---|---|---|---|---|---|---|---|---|---|---|---|---|---|---|---|---|---|---|
| | Fraction | | TBS | | TX | | FA | | TBS | | TX | | FA | | TBS | | TX | | FA | |
| Locus | SNP | A1 | β | P | β | P | β | P | β | P | β | P | β | P | β | P | β | P | β | P |
| *EPHA1* | rs10808026 | A | -0.35 | 4.31E-01 | 0.05 | 8.63E-01 | -0.03 | 6.91E-01 | -0.05 | 4.81E-01 | -1.05 | 2.90E-01 | -0.23 | 5.15E-03 | -0.05 | 4.40E-01 | 0.00 | 9.39E-01 | -0.07 | 5.07E-01 |
| *INPP5D* | rs10933431 | G | 0.00 | 9.97E-01 | 0.01 | 9.77E-01 | 0.06 | 4.15E-01 | -0.03 | 7.08E-01 | 0.26 | 7.90E-01 | 0.06 | 4.29E-01 | -0.11 | 1.03E-01 | 0.01 | 7.90E-01 | 0.07 | 4.73E-01 |
| *SORL1* | rs11218343 | C | 1.30 | 2.22E-01 | 1.53 | 2.29E-02 | -0.10 | 5.90E-01 | 0.08 | 6.48E-01 | 1.73 | 4.64E-01 | 0.11 | 5.80E-01 | 0.28 | 9.18E-02 | -0.02 | 8.24E-01 | -0.20 | 4.02E-01 |
| *OARD1* | rs114812713 | C | -0.03 | 9.76E-01 | 0.07 | 9.06E-01 | 0.16 | 2.99E-01 | -0.12 | 3.81E-01 | -2.56 | 1.91E-01 | -0.12 | 4.83E-01 | 0.02 | 8.60E-01 | 0.14 | 4.78E-02 | 0.19 | 3.29E-01 |
| *NYAP1* | rs12539172 | T | -0.09 | 8.16E-01 | 0.24 | 3.13E-01 | 0.08 | 2.24E-01 | -0.01 | 8.76E-01 | -0.14 | 8.71E-01 | -0.06 | 3.73E-01 | -0.12 | 5.38E-02 | -0.01 | 7.65E-01 | -0.03 | 7.58E-01 |
| *SLC24A4* | rs12881735 | C | -0.26 | 5.22E-01 | 0.16 | 5.35E-01 | -0.01 | 8.96E-01 | 0.02 | 8.09E-01 | 0.16 | 8.65E-01 | -0.05 | 5.18E-01 | 0.00 | 9.84E-01 | -0.02 | 5.79E-01 | -0.06 | 5.46E-01 |
| *CYB561; ACE* | rs138190086 | A | 0.59 | 6.10E-01 | -0.36 | 6.24E-01 | 0.17 | 3.96E-01 | -0.09 | 6.44E-01 | -0.88 | 7.33E-01 | 0.13 | 5.57E-01 | 0.04 | 8.40E-01 | 0.03 | 7.54E-01 | 0.13 | 6.14E-01 |
| *FERMT2* | rs17125924 | G | 0.45 | 4.57E-01 | -0.02 | 9.60E-01 | 0.03 | 7.45E-01 | 0.00 | 9.66E-01 | -1.51 | 2.65E-01 | -0.09 | 4.43E-01 | 0.00 | 9.84E-01 | 0.01 | 7.81E-01 | 0.08 | 5.61E-01 |
| *MEF2C-AS1* | rs190982 | G | 0.24 | 4.75E-01 | 0.17 | 4.31E-01 | -0.05 | 3.97E-01 | 0.08 | 1.40E-01 | 0.51 | 4.95E-01 | 0.04 | 4.99E-01 | -0.06 | 2.58E-01 | -0.03 | 2.35E-01 | 0.04 | 6.14E-01 |
| *CYYR1; ADAMTS1* | rs2830500 | A | -0.31 | 3.89E-01 | -0.41 | 7.57E-02 | -0.04 | 5.26E-01 | -0.07 | 2.10E-01 | -1.03 | 2.01E-01 | -0.09 | 1.63E-01 | -0.01 | 8.70E-01 | 0.00 | 9.09E-01 | -0.08 | 3.24E-01 |
| *SPI1* | rs3740688 | G | 0.19 | 5.76E-01 | 0.01 | 9.69E-01 | -0.07 | 2.04E-01 | 0.00 | 9.96E-01 | -0.51 | 5.09E-01 | -0.05 | 4.40E-01 | -0.05 | 3.62E-01 | 0.00 | 8.72E-01 | 0.01 | 8.62E-01 |
| *ABCA7* | rs3752246 | G | 0.32 | 4.93E-01 | 0.20 | 5.08E-01 | 0.04 | 6.06E-01 | 0.07 | 3.70E-01 | -0.50 | 6.35E-01 | -0.10 | 2.47E-01 | 0.06 | 4.43E-01 | -0.02 | 6.85E-01 | -0.04 | 7.01E-01 |
| *PICALM; EED* | rs3851179 | T | -0.27 | 4.47E-01 | -0.22 | 3.28E-01 | -0.01 | 8.75E-01 | -0.01 | 9.26E-01 | -0.44 | 5.75E-01 | 0.10 | 1.45E-01 | -0.10 | 9.09E-02 | -0.03 | 2.76E-01 | 0.05 | 5.22E-01 |
| *APOE* | rs429358 | C | -2.31 | 1.04E-10 | -0.51 | 2.82E-02 | 0.49 | 4.53E-16 | 0.02 | 7.89E-01 | -0.64 | 4.31E-01 | -0.03 | 6.32E-01 | -0.01 | 8.51E-01 | -0.02 | 6.04E-01 | 0.08 | 3.18E-01 |
| *GPR141; NME8* | rs4723711 | T | -0.28 | 4.56E-01 | -0.28 | 2.35E-01 | 0.07 | 2.64E-01 | 0.01 | 8.07E-01 | -0.12 | 8.87E-01 | 0.04 | 5.83E-01 | -0.08 | 1.60E-01 | -0.03 | 3.76E-01 | 0.01 | 9.13E-01 |
| *CR1* | rs4844610 | A | -0.63 | 1.55E-01 | -0.73 | 9.45E-03 | -0.01 | 8.73E-01 | -0.09 | 1.99E-01 | -1.15 | 2.44E-01 | 0.03 | 7.60E-01 | -0.06 | 4.15E-01 | 0.01 | 7.44E-01 | 0.05 | 6.09E-01 |
| *ADAM10;LOC101928725* | rs593742 | G | 0.08 | 8.29E-01 | 0.23 | 3.39E-01 | 0.00 | 9.56E-01 | 0.12 | 4.02E-02 | 1.41 | 9.67E-02 | -0.13 | 5.96E-02 | -0.03 | 5.97E-01 | -0.04 | 2.40E-01 | -0.18 | 3.50E-02 |
| *CASS4* | rs6024870 | A | 0.21 | 7.31E-01 | 0.01 | 9.76E-01 | 0.04 | 6.74E-01 | 0.13 | 1.88E-01 | -0.34 | 8.01E-01 | 0.08 | 4.70E-01 | -0.14 | 1.35E-01 | -0.07 | 1.35E-01 | -0.03 | 8.19E-01 |
| *WWOX; MAF* | rs62039712 | A | 0.59 | 3.61E-01 | 0.08 | 8.50E-01 | 0.08 | 4.42E-01 | 0.16 | 1.08E-01 | 2.03 | 1.59E-01 | 0.01 | 9.24E-01 | -0.14 | 1.75E-01 | -0.09 | 9.08E-02 | -0.09 | 5.27E-01 |
| *BIN1; CYP27C1* | rs6733839 | T | -0.53 | 1.11E-01 | -0.28 | 1.85E-01 | -0.06 | 2.97E-01 | -0.05 | 2.99E-01 | -0.52 | 4.88E-01 | -0.01 | 8.95E-01 | -0.01 | 8.50E-01 | 0.03 | 2.50E-01 | 0.08 | 2.73E-01 |
| *IQCK* | rs7185636 | C | 0.56 | 2.01E-01 | 0.32 | 2.48E-01 | 0.09 | 2.32E-01 | 0.05 | 4.96E-01 | 0.57 | 5.56E-01 | 0.18 | 2.73E-02 | -0.07 | 2.96E-01 | -0.03 | 3.48E-01 | 0.10 | 2.97E-01 |
| *PTK2B* | rs73223431 | T | -0.21 | 5.27E-01 | -0.10 | 6.27E-01 | -0.04 | 4.82E-01 | -0.05 | 3.37E-01 | -0.13 | 8.58E-01 | 0.01 | 8.84E-01 | -0.03 | 5.69E-01 | 0.06 | 2.43E-02 | 0.19 | 9.75E-03 |
| *USP6NL; ECHDC3* | rs7920721 | G | 0.32 | 3.40E-01 | 0.00 | 9.83E-01 | 0.00 | 9.80E-01 | 0.06 | 2.87E-01 | -0.30 | 6.87E-01 | 0.00 | 9.50E-01 | -0.06 | 3.06E-01 | -0.01 | 7.68E-01 | -0.03 | 7.30E-01 |
| *MS4A2;MS4A6A* | rs7933202 | C | 0.07 | 8.60E-01 | -0.11 | 6.57E-01 | 0.01 | 8.53E-01 | -0.01 | 8.25E-01 | -0.76 | 3.67E-01 | -0.06 | 4.26E-01 | 0.08 | 2.00E-01 | 0.00 | 9.47E-01 | -0.02 | 7.85E-01 |
| *CLU* | rs9331896 | C | 0.75 | 3.36E-02 | 0.54 | 1.63E-02 | -0.03 | 5.88E-01 | 0.12 | 3.17E-02 | 1.25 | 1.14E-01 | -0.03 | 6.27E-01 | 0.03 | 5.83E-01 | -0.07 | 1.50E-02 | -0.16 | 5.11E-02 |
| *TNFRSF21; CD2AP* | rs9473117 | C | 0.15 | 6.99E-01 | -0.46 | 5.91E-02 | 0.01 | 8.87E-01 | -0.05 | 3.97E-01 | 0.24 | 7.79E-01 | 0.03 | 6.35E-01 | -0.04 | 4.93E-01 | 0.01 | 8.43E-01 | 0.15 | 9.09E-02 |

**Table S9: Analysis of Significant AD GWAS SNPs from Kunkel et al 2019.** Results of significant SNPs identified by Kunkel et al. 2019 in their meta-analyses for each biochemical fraction. Of the 28 SNPs identified in Kunkel et al. 2019, 26 were present in our dataset and 10 were significant (p-value < 0.05) in at least one biochemical fraction. The two SNPs not present in our dataset include rs75932628 (*TREM2*) and rs9271058 (*HLA-DRB1*). Significant p-values in red text, significant positive betas are colored blue, and significant negative betas are colored red. A1= Tested allele, β= Beta value, P= P-value.

| | rsID | Proxy rsID | Taqman assay ID |
|---|---|---|---|
| **1** | rs11845003 | - | C__32007925_10 |
| **2** | rs116726862 | - | C_150768532_10 |
| **3** | rs77785770 | - | C_105058705_10 |
| **4** | rs9890231 | - | ANFVZWM |
| **5** | rs283815 | - | C_188843436_10 |
| **6** | rs116580059 | rs79606912 | C_100730255_10 |
| **7** | rs148028977 | rs146973898 | C_163754645_10 |
| **8** | rs429358 | - | C___3084793_20 |

**Table S10: Taqman Assays used for genotyping key variants.** IDs for Taqman assays used to genotype key variants or their proxies ($r^2$ = 1, D' = 1 in 1000 Genomes EUR). Variants 1- 7 were genotyped for this study, variant 8 was genotyped previously using this assay and queried from an in-house database.

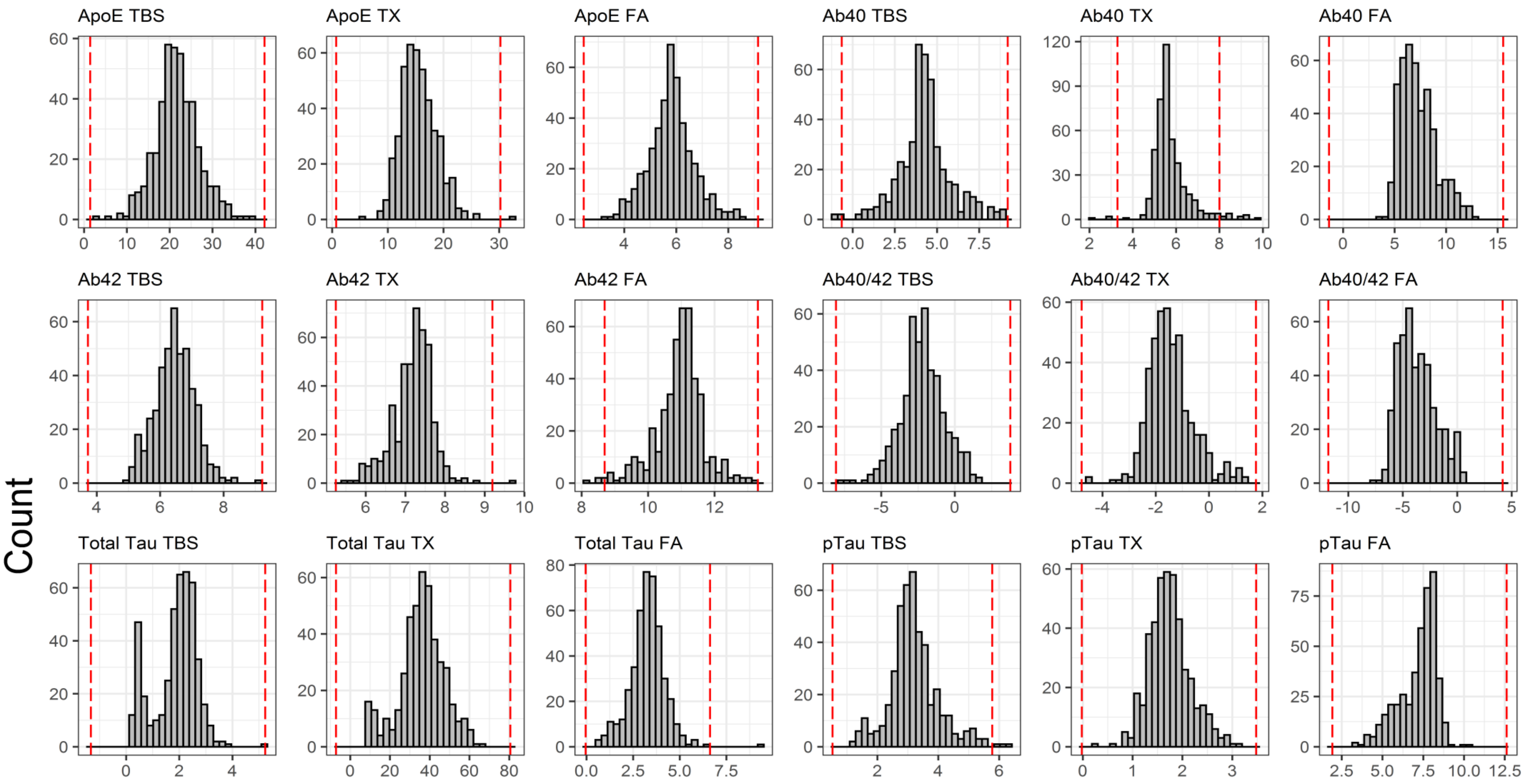


**Figure S1: Histograms of biochemical measures.** Histogram plots of all transformed biochemical measures. Dashed red lines define 3 times the interquartile range.

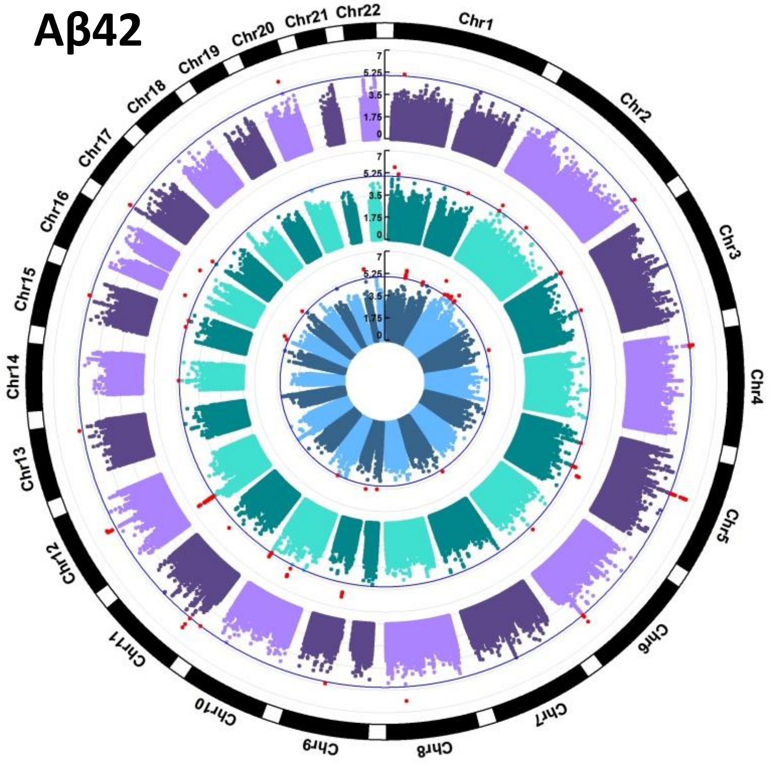


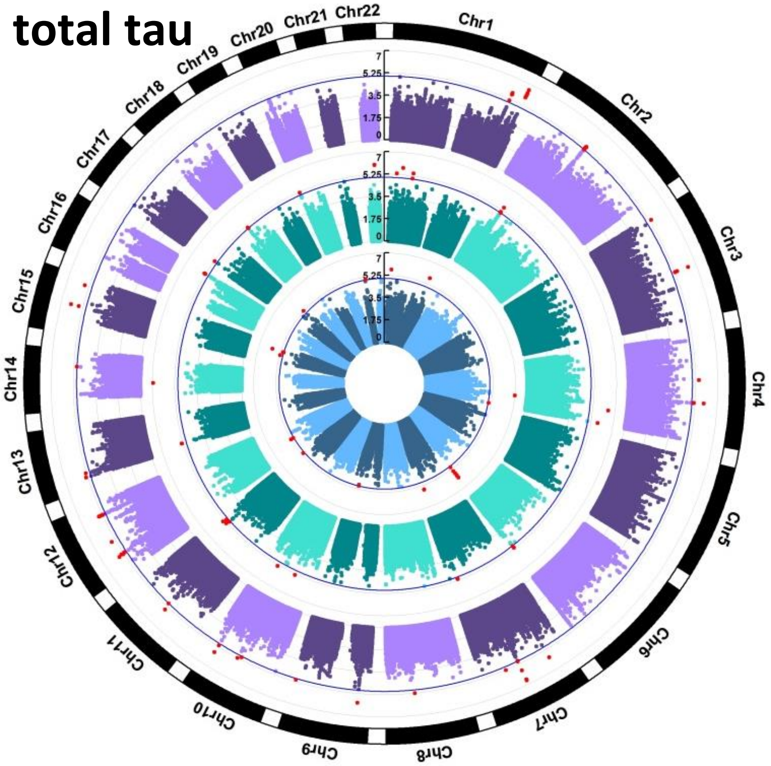


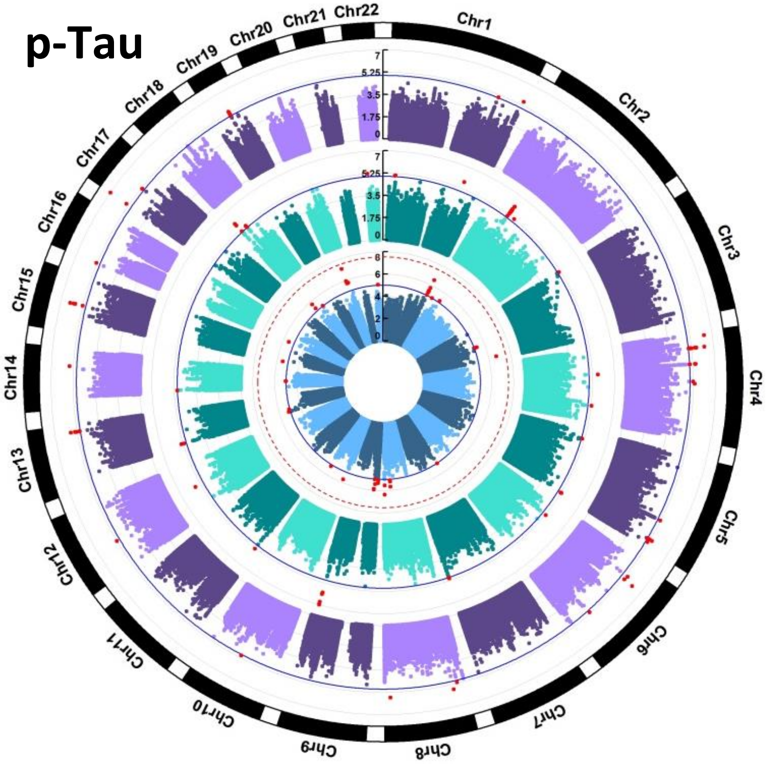


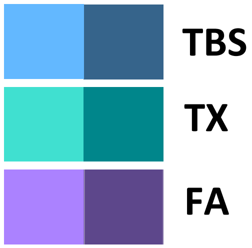


**Figure S2: Circular Manhattan Plots with no GWS SNPs.** Circular Manhattan plots for proteins and fractions with no genome-wide significance SNPs (p-value < 2.89E-08). Solid blue line marks p-value = 1E-05, red dotted line marks p-value = 2.89E-08. SNPs with a p-value < 1E-05 are colored red. Radial axes measure -log10(p-value). Inner most blue circle is the soluble TBS fraction, middle green circle is the membrane TX fraction and outer most purple circle is the insoluble FA fraction.

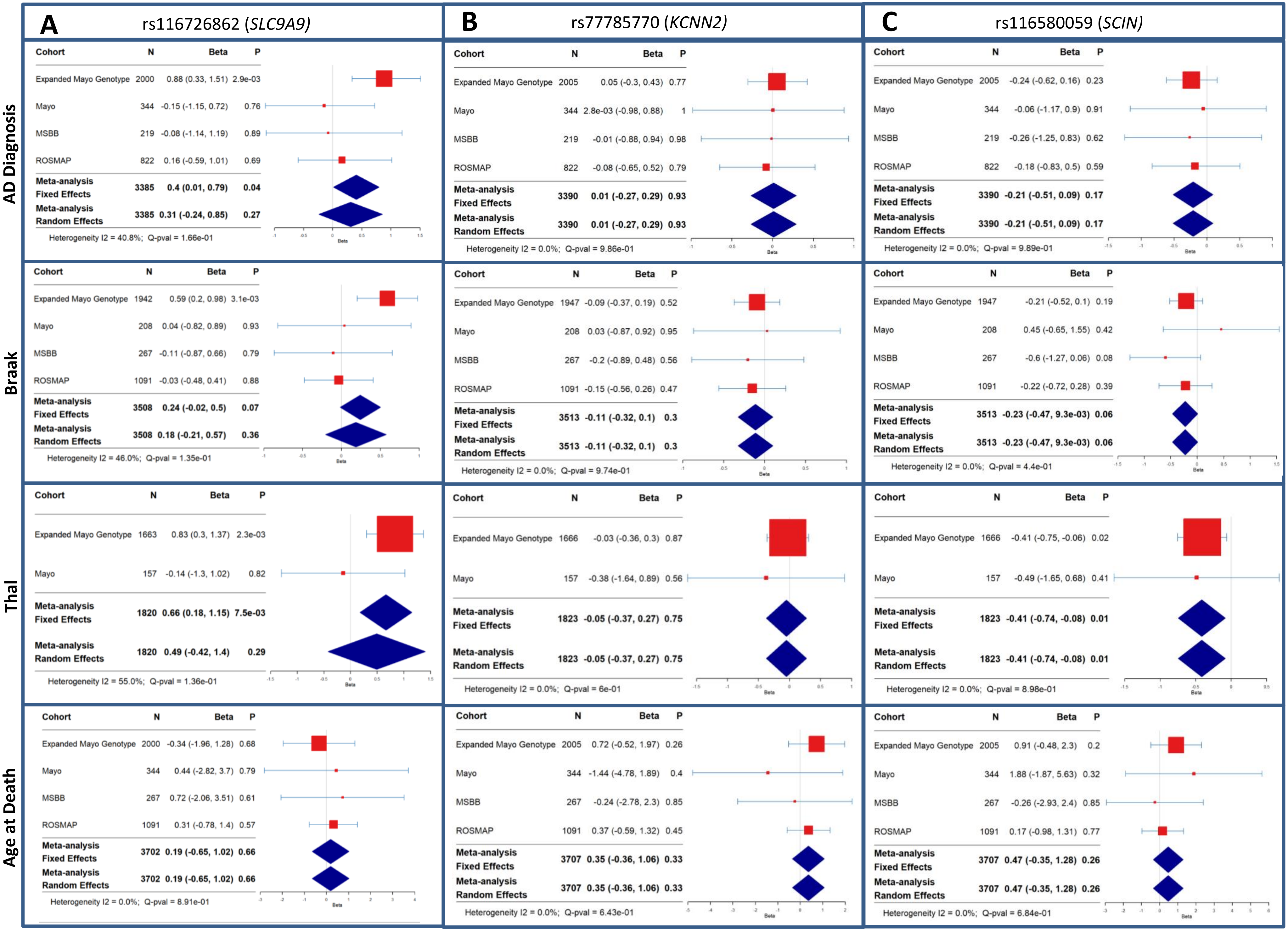


**Figure S3: Forest Plots.** Forest plots of GWS index variant associations and meta-analysis results (A-I) in independent AMP-AD and expanded Mayo genotype datasets with available AD related measures. Regression analyses include logistic for AD diagnosis, ordinal for Braak stage and Thal phase, and linear for Age at Death association tests (red boxes). Inverse variance weighted meta-analyses performed with fixed and random effects (blue diamonds).

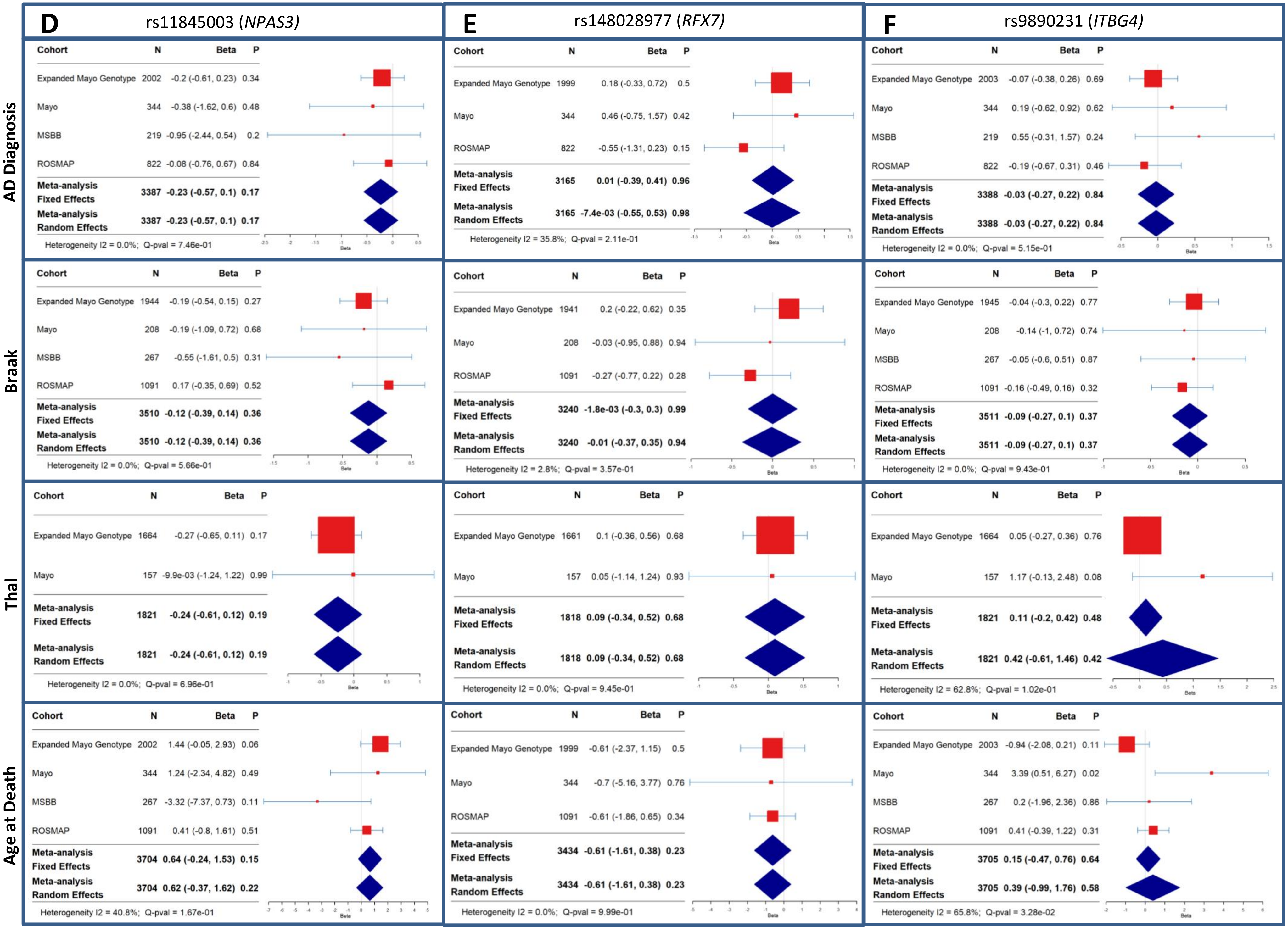
D
rs11845003 (NPAS3)
E
rs148028977 (RFX7)
F
rs9890231 (ITBG4)
AD Diagnosis
Braak
Thal
Age at Death
Cohort N Beta P
Expanded Mayo Genotype 2002 -0.2 (-0.61, 0.23) 0.34
Mayo 344 -0.38 (-1.62, 0.6) 0.48
MSBB 219 -0.95 (-2.44, 0.54) 0.2
ROSMAP 822 -0.08 (-0.76, 0.67) 0.84
Meta-analysis Fixed Effects 3387 -0.23 (-0.57, 0.1) 0.17
Meta-analysis Random Effects 3387 -0.23 (-0.57, 0.1) 0.17
Heterogeneity I2 = 0.0%; Q-pval = 7.46e-01
Cohort N Beta P
Expanded Mayo Genotype 1999 0.18 (-0.33, 0.72) 0.5
Mayo 344 0.46 (-0.75, 1.57) 0.42
ROSMAP 822 -0.55 (-1.31, 0.23) 0.15
Meta-analysis Fixed Effects 3165 0.01 (-0.39, 0.41) 0.96
Meta-analysis Random Effects 3165 -7.4e-03 (-0.55, 0.53) 0.98
Heterogeneity I2 = 35.8%; Q-pval = 2.11e-01
Cohort N Beta P
Expanded Mayo Genotype 2003 -0.07 (-0.38, 0.26) 0.69
Mayo 344 0.19 (-0.62, 0.92) 0.62
MSBB 219 0.55 (-0.31, 1.57) 0.24
ROSMAP 822 -0.19 (-0.67, 0.31) 0.46
Meta-analysis Fixed Effects 3388 -0.03 (-0.27, 0.22) 0.84
Meta-analysis Random Effects 3388 -0.03 (-0.27, 0.22) 0.84
Heterogeneity I2 = 0.0%; Q-pval = 5.15e-01
Cohort N Beta P
Expanded Mayo Genotype 1944 -0.19 (-0.54, 0.15) 0.27
Mayo 208 -0.19 (-1.09, 0.72) 0.68
MSBB 267 -0.55 (-1.61, 0.5) 0.31
ROSMAP 1091 0.17 (-0.35, 0.69) 0.52
Meta-analysis Fixed Effects 3510 -0.12 (-0.39, 0.14) 0.36
Meta-analysis Random Effects 3510 -0.12 (-0.39, 0.14) 0.36
Heterogeneity I2 = 0.0%; Q-pval = 5.66e-01
Cohort N Beta P
Expanded Mayo Genotype 1941 0.2 (-0.22, 0.62) 0.35
Mayo 208 -0.03 (-0.95, 0.88) 0.94
ROSMAP 1091 -0.27 (-0.77, 0.22) 0.28
Meta-analysis Fixed Effects 3240 -1.8e-03 (-0.3, 0.3) 0.99
Meta-analysis Random Effects 3240 -0.01 (-0.37, 0.35) 0.94
Heterogeneity I2 = 2.8%; Q-pval = 3.57e-01
Cohort N Beta P
Expanded Mayo Genotype 1945 -0.04 (-0.3, 0.22) 0.77
Mayo 208 -0.14 (-1, 0.72) 0.74
MSBB 267 -0.05 (-0.6, 0.51) 0.87
ROSMAP 1091 -0.16 (-0.49, 0.16) 0.32
Meta-analysis Fixed Effects 3511 -0.09 (-0.27, 0.1) 0.37
Meta-analysis Random Effects 3511 -0.09 (-0.27, 0.1) 0.37
Heterogeneity I2 = 0.0%; Q-pval = 9.43e-01
Cohort N Beta P
Expanded Mayo Genotype 1664 -0.27 (-0.65, 0.11) 0.17
Mayo 157 -9.9e-03 (-1.24, 1.22) 0.99
Meta-analysis Fixed Effects 1821 -0.24 (-0.61, 0.12) 0.19
Meta-analysis Random Effects 1821 -0.24 (-0.61, 0.12) 0.19
Heterogeneity I2 = 0.0%; Q-pval = 6.96e-01
Cohort N Beta P
Expanded Mayo Genotype 1661 0.1 (-0.36, 0.56) 0.68
Mayo 157 0.05 (-1.14, 1.24) 0.93
Meta-analysis Fixed Effects 1818 0.09 (-0.34, 0.52) 0.68
Meta-analysis Random Effects 1818 0.09 (-0.34, 0.52) 0.68
Heterogeneity I2 = 0.0%; Q-pval = 9.45e-01
Cohort N Beta P
Expanded Mayo Genotype 1664 0.05 (-0.27, 0.36) 0.76
Mayo 157 1.17 (-0.13, 2.48) 0.08
Meta-analysis Fixed Effects 1821 0.11 (-0.2, 0.42) 0.48
Meta-analysis Random Effects 1821 0.42 (-0.61, 1.46) 0.42
Heterogeneity I2 = 62.8%; Q-pval = 1.02e-01
Cohort N Beta P
Expanded Mayo Genotype 2002 1.44 (-0.05, 2.93) 0.06
Mayo 344 1.24 (-2.34, 4.82) 0.49
MSBB 267 -3.32 (-7.37, 0.73) 0.11
ROSMAP 1091 0.41 (-0.8, 1.61) 0.51
Meta-analysis Fixed Effects 3704 0.64 (-0.24, 1.53) 0.15
Meta-analysis Random Effects 3704 0.62 (-0.37, 1.62) 0.22
Heterogeneity I2 = 40.8%; Q-pval = 1.67e-01
Cohort N Beta P
Expanded Mayo Genotype 1999 -0.61 (-2.37, 1.15) 0.5
Mayo 344 -0.7 (-5.16, 3.77) 0.76
ROSMAP 1091 -0.61 (-1.86, 0.65) 0.34
Meta-analysis Fixed Effects 3434 -0.61 (-1.61, 0.38) 0.23
Meta-analysis Random Effects 3434 -0.61 (-1.61, 0.38) 0.23
Heterogeneity I2 = 0.0%; Q-pval = 9.99e-01
Cohort N Beta P
Expanded Mayo Genotype 2003 -0.94 (-2.08, 0.21) 0.11
Mayo 344 3.39 (0.51, 6.27) 0.02
MSBB 267 0.2 (-1.96, 2.36) 0.86
ROSMAP 1091 0.41 (-0.39, 1.22) 0.31
Meta-analysis Fixed Effects 3705 0.15 (-0.47, 0.76) 0.64
Meta-analysis Random Effects 3705 0.39 (-0.99, 1.76) 0.58
Heterogeneity I2 = 65.8%; Q-pval = 3.28e-02
Beta

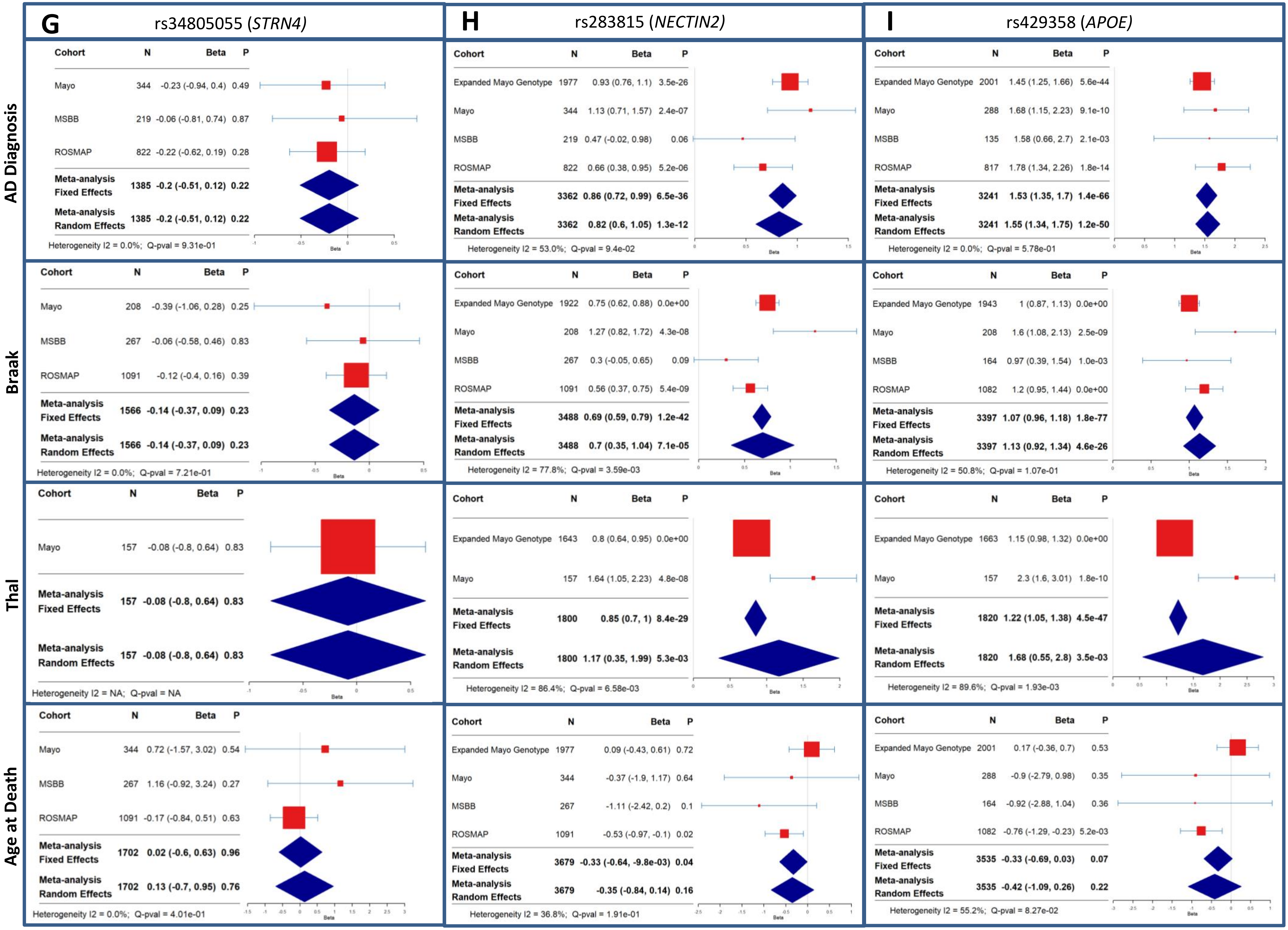
G
rs34805055 (STRN4)
H
rs283815 (NECTIN2)
I
rs429358 (APOE)
AD Diagnosis
Braak
Thal
Age at Death
Cohort N Beta P
Mayo 344 -0.23 (-0.94, 0.4) 0.49
MSBB 219 -0.06 (-0.81, 0.74) 0.87
ROSMAP 822 -0.22 (-0.62, 0.19) 0.28
Meta-analysis Fixed Effects 1385 -0.2 (-0.51, 0.12) 0.22
Meta-analysis Random Effects 1385 -0.2 (-0.51, 0.12) 0.22
Heterogeneity I2 = 0.0%; Q-pval = 9.31e-01
Cohort N Beta P
Expanded Mayo Genotype 1977 0.93 (0.76, 1.1) 3.5e-26
Mayo 344 1.13 (0.71, 1.57) 2.4e-07
MSBB 219 0.47 (-0.02, 0.98) 0.06
ROSMAP 822 0.66 (0.38, 0.95) 5.2e-06
Meta-analysis Fixed Effects 3362 0.86 (0.72, 0.99) 6.5e-36
Meta-analysis Random Effects 3362 0.82 (0.6, 1.05) 1.3e-12
Heterogeneity I2 = 53.0%; Q-pval = 9.4e-02
Cohort N Beta P
Expanded Mayo Genotype 2001 1.45 (1.25, 1.66) 5.6e-44
Mayo 288 1.68 (1.15, 2.23) 9.1e-10
MSBB 135 1.58 (0.66, 2.7) 2.1e-03
ROSMAP 817 1.78 (1.34, 2.26) 1.8e-14
Meta-analysis Fixed Effects 3241 1.53 (1.35, 1.7) 1.4e-66
Meta-analysis Random Effects 3241 1.55 (1.34, 1.75) 1.2e-50
Heterogeneity I2 = 0.0%; Q-pval = 5.78e-01
Cohort N Beta P
Mayo 208 -0.39 (-1.06, 0.28) 0.25
MSBB 267 -0.06 (-0.58, 0.46) 0.83
ROSMAP 1091 -0.12 (-0.4, 0.16) 0.39
Meta-analysis Fixed Effects 1566 -0.14 (-0.37, 0.09) 0.23
Meta-analysis Random Effects 1566 -0.14 (-0.37, 0.09) 0.23
Heterogeneity I2 = 0.0%; Q-pval = 7.21e-01
Cohort N Beta P
Expanded Mayo Genotype 1922 0.75 (0.62, 0.88) 0.0e+00
Mayo 208 1.27 (0.82, 1.72) 4.3e-08
MSBB 267 0.3 (-0.05, 0.65) 0.09
ROSMAP 1091 0.56 (0.37, 0.75) 5.4e-09
Meta-analysis Fixed Effects 3488 0.69 (0.59, 0.79) 1.2e-42
Meta-analysis Random Effects 3488 0.7 (0.35, 1.04) 7.1e-05
Heterogeneity I2 = 77.8%; Q-pval = 3.59e-03
Cohort N Beta P
Expanded Mayo Genotype 1943 1 (0.87, 1.13) 0.0e+00
Mayo 208 1.6 (1.08, 2.13) 2.5e-09
MSBB 164 0.97 (0.39, 1.54) 1.0e-03
ROSMAP 1082 1.2 (0.95, 1.44) 0.0e+00
Meta-analysis Fixed Effects 3397 1.07 (0.96, 1.18) 1.8e-77
Meta-analysis Random Effects 3397 1.13 (0.92, 1.34) 4.6e-26
Heterogeneity I2 = 50.8%; Q-pval = 1.07e-01
Cohort N Beta P
Mayo 157 -0.08 (-0.8, 0.64) 0.83
Meta-analysis Fixed Effects 157 -0.08 (-0.8, 0.64) 0.83
Meta-analysis Random Effects 157 -0.08 (-0.8, 0.64) 0.83
Heterogeneity I2 = NA; Q-pval = NA
Cohort N Beta P
Expanded Mayo Genotype 1643 0.8 (0.64, 0.95) 0.0e+00
Mayo 157 1.64 (1.05, 2.23) 4.8e-08
Meta-analysis Fixed Effects 1800 0.85 (0.7, 1) 8.4e-29
Meta-analysis Random Effects 1800 1.17 (0.35, 1.99) 5.3e-03
Heterogeneity I2 = 86.4%; Q-pval = 6.58e-03
Cohort N Beta P
Expanded Mayo Genotype 1663 1.15 (0.98, 1.32) 0.0e+00
Mayo 157 2.3 (1.6, 3.01) 1.8e-10
Meta-analysis Fixed Effects 1820 1.22 (1.05, 1.38) 4.5e-47
Meta-analysis Random Effects 1820 1.68 (0.55, 2.8) 3.5e-03
Heterogeneity I2 = 89.6%; Q-pval = 1.93e-03
Cohort N Beta P
Mayo 344 0.72 (-1.57, 3.02) 0.54
MSBB 267 1.16 (-0.92, 3.24) 0.27
ROSMAP 1091 -0.17 (-0.84, 0.51) 0.63
Meta-analysis Fixed Effects 1702 0.02 (-0.6, 0.63) 0.96
Meta-analysis Random Effects 1702 0.13 (-0.7, 0.95) 0.76
Heterogeneity I2 = 0.0%; Q-pval = 4.01e-01
Cohort N Beta P
Expanded Mayo Genotype 1977 0.09 (-0.43, 0.61) 0.72
Mayo 344 -0.37 (-1.9, 1.17) 0.64
MSBB 267 -1.11 (-2.42, 0.2) 0.1
ROSMAP 1091 -0.53 (-0.97, -0.1) 0.02
Meta-analysis Fixed Effects 3679 -0.33 (-0.64, -9.8e-03) 0.04
Meta-analysis Random Effects 3679 -0.35 (-0.84, 0.14) 0.16
Heterogeneity I2 = 36.8%; Q-pval = 1.91e-01
Cohort N Beta P
Expanded Mayo Genotype 2001 0.17 (-0.36, 0.7) 0.53
Mayo 288 -0.9 (-2.79, 0.98) 0.35
MSBB 164 -0.92 (-2.88, 1.04) 0.36
ROSMAP 1082 -0.76 (-1.29, -0.23) 5.2e-03
Meta-analysis Fixed Effects 3535 -0.33 (-0.69, 0.03) 0.07
Meta-analysis Random Effects 3535 -0.42 (-1.09, 0.26) 0.22
Heterogeneity I2 = 55.2%; Q-pval = 8.27e-02
Beta

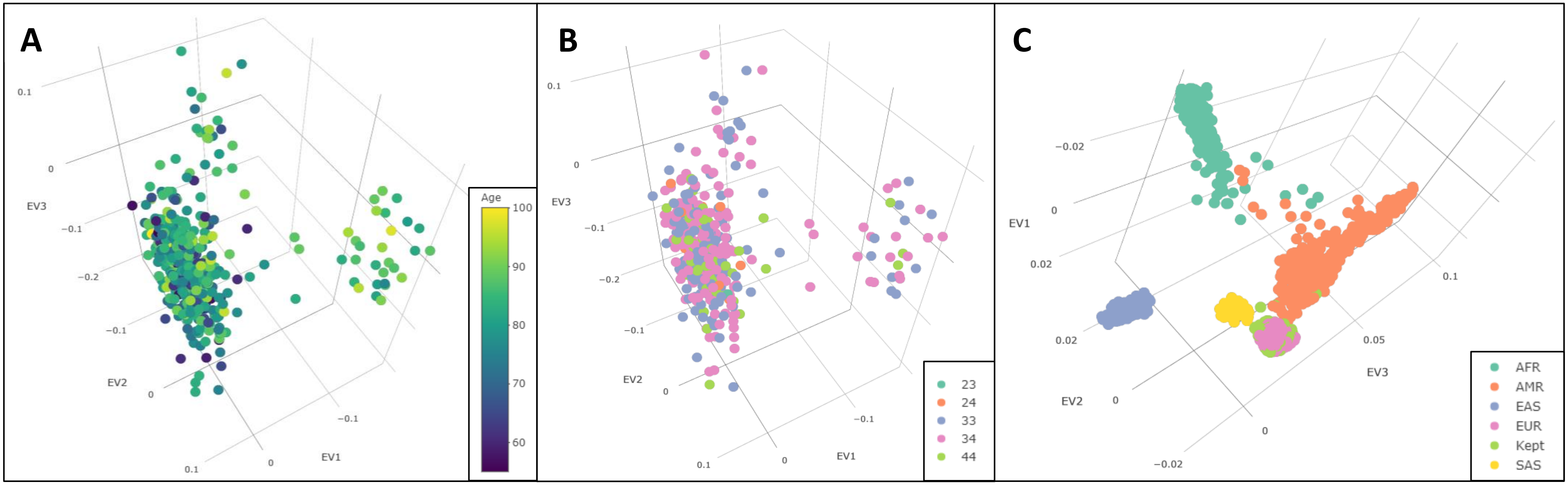


**Figure S4: Population Substructure via Eigenstrat Analysis.** Screenshots of the first three eigenvectors (EV1-3) plotted from the parent population (n=460) of our study cohort after analysis by Eigenstrat. **A:** Parent population colored by age at death. **B:** Parent population colored by *APOE* genotype. **C:** Superimposed view of study population (light green) with 1000 genomes populations.

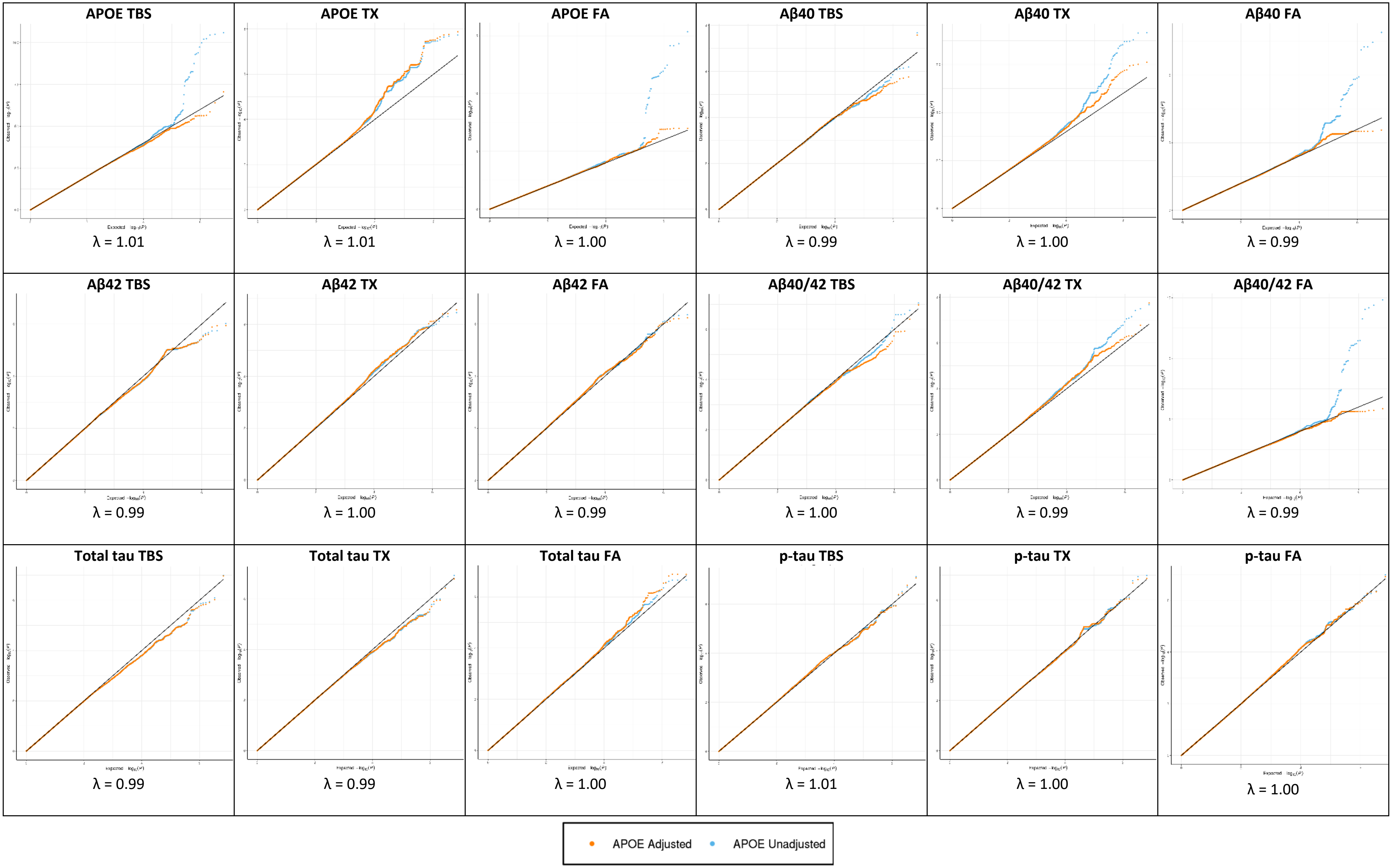


**Figure S5: Quantile-Quantile (QQ) Plots of biochemical measure GWAS.** QQ plots of each biochemical measure in *APOE-ε*2 and *APOE-ε*4 unadjusted (blue) and adjusted (orange) models. Genomic inflation values (λ) for *APOE* unadjusted models at the bottom of each plot. x-axis= expected –log10(p-value), y-axis = observed –log10(p-value).
